# Transcriptomic and proteomic retinal pigment epithelium signatures of age-related macular degeneration

**DOI:** 10.1101/2021.08.19.457044

**Authors:** Anne Senabouth, Maciej Daniszewski, Grace E. Lidgerwood, Helena H. Liang, Damián Hernández, Mehdi Mirzaei, Ran Zhang, Xikun Han, Drew Neavin, Louise Rooney, Isabel Lopez Sanchez, Lerna Gulluyan, Joao A Paulo, Linda Clarke, Lisa S Kearns, Vikkitharan Gnanasambandapillai, Chia-Ling Chan, Uyen Nguyen, Angela M Steinmann, Rachael Zekanovic, Nona Farbehi, Vivek K. Gupta, David A Mackey, Guy Bylsma, Nitin Verma, Stuart MacGregor, Robyn H Guymer, Joseph E. Powell, Alex W. Hewitt, Alice Pébay

**Author notes:** Equal first authors (Anne Senabouth, Maciej Daniszewski, Grace E. Lidgerwood). Equal senior authors (Joseph E. Powell, Alex W. Hewitt, Alice Pébay).

## Abstract

Induced pluripotent stem cells generated from patients with geographic atrophy as well as healthy individuals were differentiated to retinal pigment epithelium (RPE) cells. By integrating transcriptional profiles of 127,659 RPE cells generated from 43 individuals with geographic atrophy and 36 controls with genotype data, we identified 439 expression Quantitative Trait (eQTL) loci in cis that were associated with disease status and specific to subpopulations of RPE cells. We identified loci linked to two genes with known associations with geographic atrophy - PILRB and PRPH2, in addition to 43 genes with significant genotype x disease interactions that are candidates for novel genetic associations for geographic atrophy. On a transcriptome-only level, we identified molecular pathways significantly upregulated in geographic atrophy-RPE including in extracellular cellular matrix reorganisation, neurodegeneration, and mitochondrial functions. We subsequently implemented a large-scale proteomics analysis, confirming modification in proteins associated with these pathways. We also identified six significant protein (p) QTL that regulate protein expression in the RPE cells and in geographic atrophy - two of which share variants with cis-eQTL. Transcriptome-wide association analysis identified genes at loci previously associated with age-related macular degeneration. Further analysis conditional on disease status, implicated statistically significant RPE-specific eQTL. This study uncovers important differences in RPE homeostasis associated with geographic atrophy.

---

Age-related macular degeneration (AMD) is a progressive, degenerative disease caused by dysfunction and death of the retinal pigment epithelium (RPE), and photoreceptors, leading to irreversible vision loss. AMD is the leading cause of vision loss and legal blindness in higher resourced countries ^1^. There are two forms of the vision threatening late stage of AMD; neovascular and geographic atrophy ^2^, the latter affecting more than 5 million people globally ^3^. Whilst management of neovascular AMD has improved significantly since the introduction of intravitreal anti-vascular endothelial growth factor (VEGF) injections ^4–7^, there are currently no approved or effective treatments for geographic atrophy, despite multiple clinical trials to evaluate potential drug candidates and interventions ^8–13^. This presents a significant unmet medical need and as such, greater effort in disease modelling and drug discovery should be aimed at preventing and delaying disease progression.

It is now well established that both environmental and genetic risk factors contribute to AMD ^14^. A common variant in the *CFH* gene (*CFH* Y402H) is estimated to account for nearly half of all AMD risk ^15–18^. Furthermore, variants at the *LOC387715/ARMS2/HTRA1* locus have been identified as major contributors to AMD development ^19, 20^. To date, genome-wide association studies (GWAS) have identified over 30 independent loci where a common risk allele is associated with an increased risk of AMD ^21–23^. These loci influence distinct biological pathways, including the complement system, lipid transport, extracellular matrix remodelling, angiogenesis and cell survival ^24^.

Unlike rare and highly penetrant variants that largely contribute to disease by altering protein sequences, common variants predominantly act via changes in gene regulation ^25^. Mapping expression quantitative trait loci (eQTL) is a powerful approach to elucidate functional mechanisms of common genetic variants, allowing the allelic effect of a variant on disease risk to be linked to changes in gene expression. Three recent studies applied eQTL mapping in post-mortem retina to investigate the regulation of gene expression and identified eQTL variants regulating gene expression with a subset of these eQTL associated with AMD in GWAS ^26–28^. Molecular and genetic profiling of RPE in healthy and diseased tissue would likely improve our understanding of the mechanisms that confer disease risk or contribute to geographic atrophy progression. However, the invasive nature of retina harvest highly restricts tissue availability to post-mortem donors. This limitation can be overcome by reprogramming somatic cells from affected patients into patient-specific induced pluripotent stem cells (iPSCs) ^29, 30^ and subsequently differentiate them into homogenous RPE cultures for downstream disease modelling. Here, we used scRNA-seq and mass spectrometry to characterize the transcriptomic and proteomic profiles of RPE cells generated from a large cohort of iPSCs derived from healthy and geographic atrophy patients.

## Results

### Generation of patient iPSCs, differentiation to RPE cells and genomic profiling

We reprogrammed fibroblasts into iPSCs from 63 individuals with geographic atrophy (all of Northern European descent of whom 37 were female; mean ± SD age at recruitment: 83.8 ± 8.2 years) using episomal vectors as we described ^31^, with lines from 47 individuals successfully reprogrammed (**Figures S1, S2**). We matched these iPSCs with control iPSC lines from ethnically-matched healthy individuals that were generated and characterised in a previous study^32^ (**Figures S1, S2**, **Supplementary Data 1**). Lines were genotyped for 787,443 single nucleotide polymorphisms (SNPs) and imputed with the Haplotype Reference Consortium panel ^33^. After quality control, this yielded 4,309,001 autosomal SNPs with minor allele frequency (MAF) >10%. The differentiation of all iPSC lines to RPE was performed in two large independent differentiation batches, and lines that did not differentiate sufficiently to RPE were discarded from analysis (Figures 1a**, S1, S2**). Differentiated cell lines were divided into 12 pools that each consisted of up to 8 cell lines from both control and AMD groups. scRNA-seq was performed on all pools, with the targeted capture of 20,000 cells per pool and sequencing depth of 30,000 reads per cell (**Table S1**). Resulting single cell transcriptome profiles then underwent quality control and donor assignment. 18,820 cells were designated doublets and removed from the dataset, in addition to cells from individuals that were removed from the study due to low number of assigned cells (4), failed virtual karyotype (1) and failed genotype (4) (**Figure S2, Table S2**). A total of 127,659 cells from 79 individual lines remained following quality control. These include 43 geographic atrophy lines (73,161 cells, 15 males, 28 females, 83.4 ± 8.6 years) and 36 control lines (54,498 cells, 19 males, 17 females, mean ± SD age of samples 67.6 ± 9.5 years) (**Figure S2, Supplementary Data 1**).

**Figure 1.**
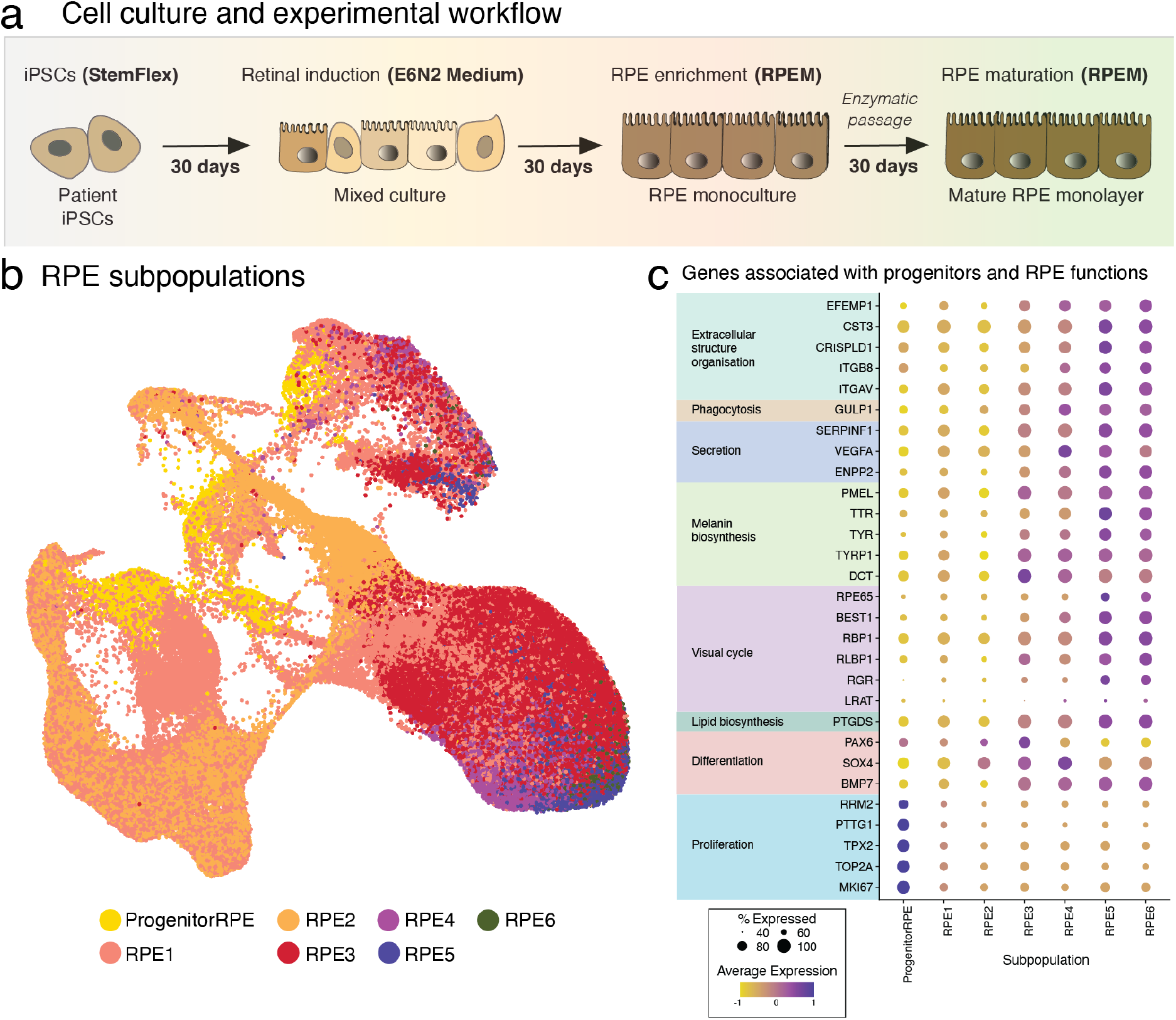
Generation of RPE from iPSCs, identification and characterisation of RPE subpopulations. **(a)** Schematic representation of the cell culture flow, with iPSCs differentiated into RPE cells in 90 days prior to harvest for scRNA-seq and mass spectrometry. **(b)** Uniform Manifold Approximation and Projection (UMAP) of cells labelled by subpopulation. Cells were assigned to subpopulations identified in a previous study using a supervised classification method, and coloured by subpopulation. **(c)** Dotplot representation of average expression of genes associated with RPE functions (extracellular structure organisation, phagocytosis, secretion, melanin biosynthesis, visual cycle, lipid biosynthesis, differentiation and proliferation) and progenitors (differentiation, proliferation) in the various subpopulations. Levels of gene expression per cell are shown with colour gradients, and frequencies of cells expressing the respective gene (% expressed) are shown with size of dots.

**Table 1.** Summary of cells retrieved in each RPE subpopulation.

| **Subpopulation** | **Number of cells (Control)** | **Number of cells (GA)** | **Total number of cells** | **% of Control cells** | **% of GA cells** |
| --- | --- | --- | --- | --- | --- |
| Progenitor RPE | 3,159 | 2,977 | 6,136 | 5.8% | 4.1% |
| RPE1 | 24,027 | 32,254 | 56,281 | 44.1% | 44.1% |
| RPE2 | 16,780 | 14,620 | 31,400 | 30.8% | 19.9% |
| RPE3 | 5,851 | 14,501 | 20,352 | 10.7% | 19.8% |
| RPE4 | 2,113 | 4,329 | 6,442 | 3.9% | 5.9% |
| RPE5 | 1,396 | 2,269 | 3,665 | 2.6% | 3.1% |
| RPE6 | 1,172 | 2,211 | 3,383 | 2.1% | 3.0% |
| Total | 54,498 | 73,161 | 127,659 | 100.0% | 100.0% |

Abbreviation: RPE, retinal pigmented epithelium; GA, geographic atrophy.

**Table 2.** Summary of Differentially-Expressed Genes in Geographic Atrophy (GA)

| Subpopulation | Upregulated | Downregulated | Total | Known Association with GA |
| --- | --- | --- | --- | --- |
| Progenitor RPE | 54 | 131 | 185 | 2 |
| RPE1 | 1,987 | 578 | 2,565 | 22 |
| RPE2 | 1,361 | 328 | 1,689 | 15 |
| RPE3 | 156 | 88 | 244 | 4 |
| RPE4 | 126 | 15 | 141 | 5 |
| RPE5 | 36 | 68 | 104 | 3 |
| RPE6 | 41 | 43 | 84 | 1 |

### Identification of seven RPE subpopulations using supervised classification

We previously used scRNA-seq to analyze the transcriptomic signature of human embryonic stem cell-derived RPE cells over 12 months in culture and identified 17 RPE subpopulations of varying levels of maturity^34^. We used this resource to build a prediction model for scPred, a supervised classification method^35^. We calculated the probabilities of each cell belonging to a reference subpopulation, and cells were assigned to the reference subpopulation with the greatest probability. While all 17 reference subpopulations were detected in this dataset, five subpopulations had fewer than 20 cells (**Table S3**). Cells from these subpopulations were excluded from further analysis, in addition to cells from donors with fewer than 20 cells in a subpopulation. This left 127,659 cells (54,498 control, 73,161 geographic atrophy cells) distributed among the remaining 7 subpopulations, with cells being classified as “RPE progenitors” (Progenitor RPE) and RPE cells (RPE1-6) (**Tables 1,** Figure 1). A Chi-Squared Test of Independence observed statistically significant differences in the proportions of subpopulations between cases and controls (*X*^2^ (6, N = 127,659) = 3672.4, p < 2.2 × 10^-16^, Table 1). Post-hoc pairwise comparisons revealed that the proportion of all subpopulations except RPE1 differed between cases and controls (**Table S4)**, and there was also variation in the proportions of subpopulations between individual cell lines (**Table S5, Figure S3**).

Genes associated with cell proliferation (*MKI67*, *TOP2A*, *TPX2, PTTG1*, *RRM2*), expressed in progenitors and differentiating RPE cells ^34^ were most highly expressed by cells of the “Progenitor RPE” and RPE1 subpopulations, indicative of a differentiating and immature RPE phenotype. The high expression of the early retinal development marker *PAX6* in RPE2 also suggests an early RPE stage within this population (Figure 1c). RPE markers were observed including genes associated with extracellular structure organization (*CST3*, *EFEMP1*, *ITGAV*, *CRISPLD1*, *ITGB8*), phagocytosis (*GULP1*), secretion (*SERPINF1*, *VEGFA*, *ENPP2*), secretion melanin biosynthesis (*PMEL*, *TTR*, *TYR*, *TYRP1*, *DCT*), visual cycle (*RPE65*, *BEST1*, *RBP1*, *RLBP1*, *RGR, LRAT*), and lipid biosynthesis (*PTGDS*) (Figure 1c). The RPE genes *PMEL*, *TYR* and *RBP1* were most highly expressed in the subpopulations RPE2-6, whilst *RGR* and *RPE65* were mainly expressed in RPE3, RPE5 and RPE6 (Figure 1c). Other genes commonly expressed in native RPE cells such as *ITGB8*, *EFEMP1*, *ITGAV*, *GULP1*, *RLBP1*, *RBP1*, *LRAT* were also enriched in RPE2-6 (Figure 1c).

### RPE subpopulations diverge into two trajectories

We used trajectory inference to identify the global lineage structure of all cells and subsequently the developmental trajectories of RPE subpopulations, using the most immature subpopulation - Progenitor RPE, as the origin. We observed a bifurcating trajectory that diverged at RPE3 to form two branches that terminate with RPE6 (Trajectory 1) and RPE4 (Trajectory 2) (Figure 2a). We applied trajectory-based differential expression analysis^36^ and observed the transition from progenitors to RPE was driven by 1,353 genes mainly involved in cell cycle (*CDKN1C, CENPK, CRABP1, DUSP1, NBL1, PRC1, RELN*); differentiation (*CRABP1, DLK1, FAM161A, NRP2, OLFM1, PCSK1N, PLXNA4*); cytoskeleton, adhesion and migration (ARPC5, *ERMN*, *ITGB1*), various metabolic processes (*SAT1, SLC16A8*, *SLC7A5*), stress (*MGST1, SGK1*), calcium transport and homeostasis (*ATP2B2, STC2*), melanin biogenesis (*TYR, TYRP1*) or translation (*EIF4EBP1*) (Figure 2b). It is not surprising that these genes show a temporary expression pattern, which would coincide with a differentiation process from a progenitor cell to a differentiating and differentiated RPE cell. Bifurcation of the trajectories was driven by 26 genes that were enriched for mitotic processes (Figure 2c). Genes driving the resulting two trajectories were very similar, with trajectory 1 and trajectory 2 sharing 99.4% of genes (**Supplementary Data 2**). For instance, trajectory 1 included genes involved in ECM organisation (*TSPAN8*), melanogenesis (*TPH1*) or retinal development (*IRX6*); and trajectory 2 expressed genes associated with lipid metabolism (*ADIRF*, *APOA1*, *CD36*), iron binding (*LCN2*, *MT1G*), cytoskeleton (*MYL7*) or retinal development (*PITX3*). Those variations do not point to clear differences in the two trajectories. Instead, these subtle and rare differences (31 genes) suggest a close resemblance of the two trajectories and further confirm the efficacy of the differentiation protocol in generating homogenous populations of RPE cells in which variations between cohorts could be attributed to the disease status rather than variabilities of differentiation.

**Figure 2.**
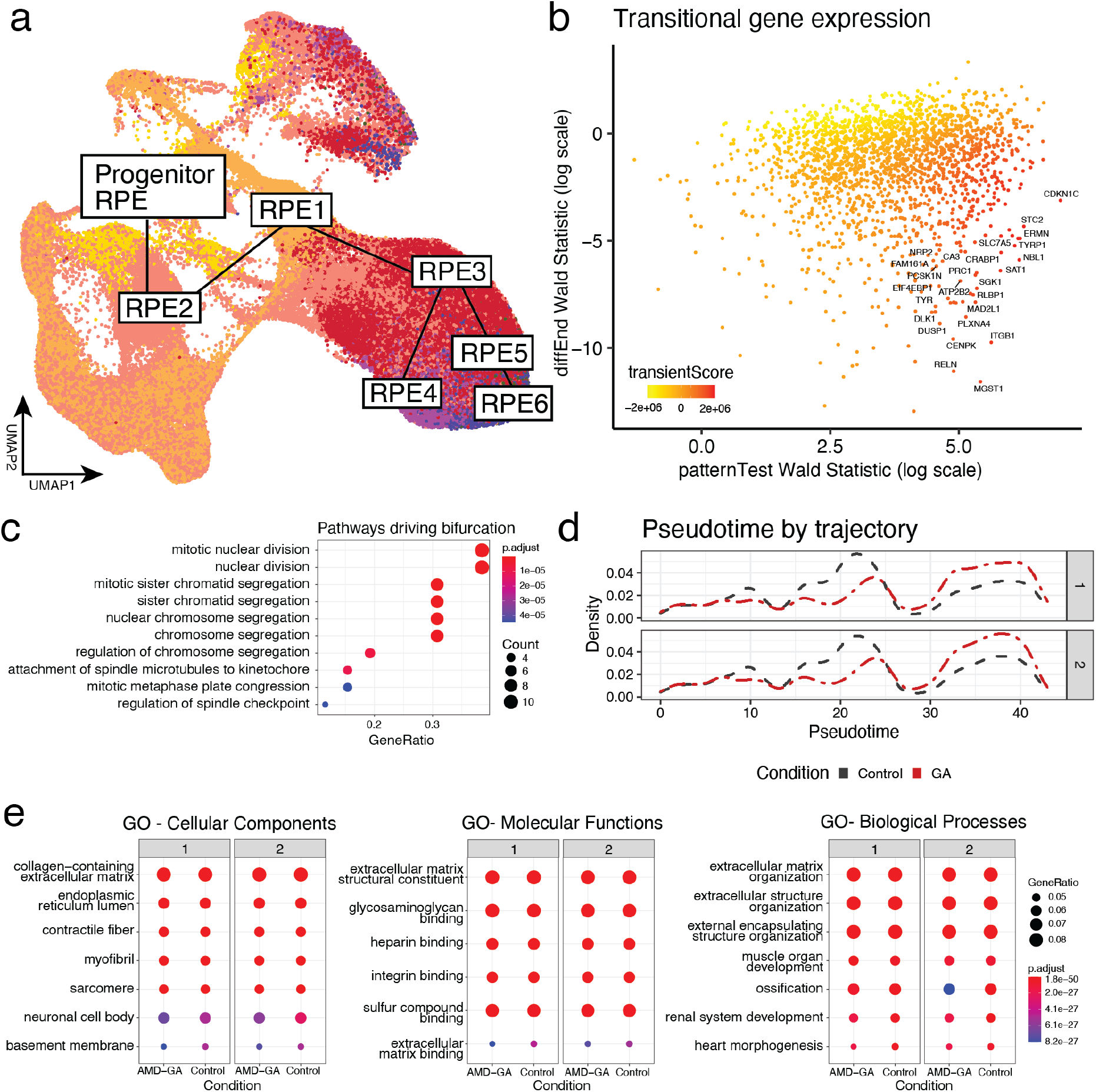
Trajectory analysis of the various subpopulations. **(a)** Uniform Manifold Approximation and Projection (UMAP) of cells labelled by subpopulation showing the two developmental trajectories bifurcating at RPE3. **(b)** Identification of transiently-expressed genes by contrasting rankings of genes (transientScore) that are differentially expressed during trajectory progression (patternTest) against expression of genes at trajectory end points (diffEnd). **(c)** Enrichment of genes that are differentially expressed during the bifurcation of the main trajectory. **(d)** Density of cells across pseudotime. Grey dashed lines represent density of cells from control donors, while red dashed lines represent density of cells from AMD-GA lines. **(e)** Enrichment of genes differentially expressed across pseudotime, that are associated with trajectories and conditions.

To determine if lineages differed based on disease status, we first tested whether the distribution of cells based within each condition differed across pseudotime - a measure of progression through the global trajectory, and noted a difference (Kolmogorov–Smirnov test. Trajectory 1 - *p*-value < 2.2 x10^-16^; Trajectory 2 - *p*-value < 2.2 x10^-16^; Figure 2d). We then assessed differential expression patterns across the whole trajectory based on disease status and after Bonferroni correction, identified 91 genes that were significant (**Supplementary Data 2**). Seven of these genes - *C3*^37^*, CRYAB*^38^*, IL6*^39^*, IL8/CXCL8*^39^, *EFEMP1*^40^, *GFAP*^41^, and *TRPM3*^21^, have previously been linked to geographic atrophy and were differentially expressed in both trajectories between control and geographic atrophy. Altogether, the differences of gene expression observed between the two cohorts likely reflect subtle differentiation differences and subsequent characteristics between RPE cells of healthy individuals and those prone to develop geographic atrophy.

### Geographic atrophy-RPE cells show specific differential gene expressions

Next, we identified genes associated with disease status in each RPE subpopulation using differential gene expression analysis (DGE), disease ontology (DO) and over-representation analysis (ORA). We identified 5,012 events of differential expression, consisting of 3,240 genes that were either upregulated or downregulated in geographic atrophy subpopulations compared to controls (**Supplementary Data 3**). The majority of differentially expressed genes were found in the two largest subpopulations - RPE1 (2,565 genes) and RPE2 (1,689 genes) (Table 2), and most genes were solely differentially expressed in a single subpopulation (Figure 3a). We identified 27 genes with known associations with Geographic Atrophy and six genes with known associations with neovascular AMD (retrieved using disGeNET v7^42^), such as *PNPLA2*, *MFGE8*, *SERPINF1*, *C3*, *VEGFA*, *HTRA1*, *CFH*, *VIM*, *STK19*, *CRYAB*, *CFI*, *CNN2*, *LRPAP1*, *RDH5*, *IMDH1*, *CFD*, *CFHR1*, *TSPO*, *APOE* and *EFEMP1* (Figure 3b). Disease ontology analysis of these genes revealed association with multiple diseases including macular degeneration, retinal degeneration, diabetic retinopathy and retinal vascular disease, Alzheimer’s disease and tauopathy, vitiligo, metabolism disorders and various cancers - as annotated by the Disease Ontology database ^43^ (**Figure S3b**).

**Figure 3.**
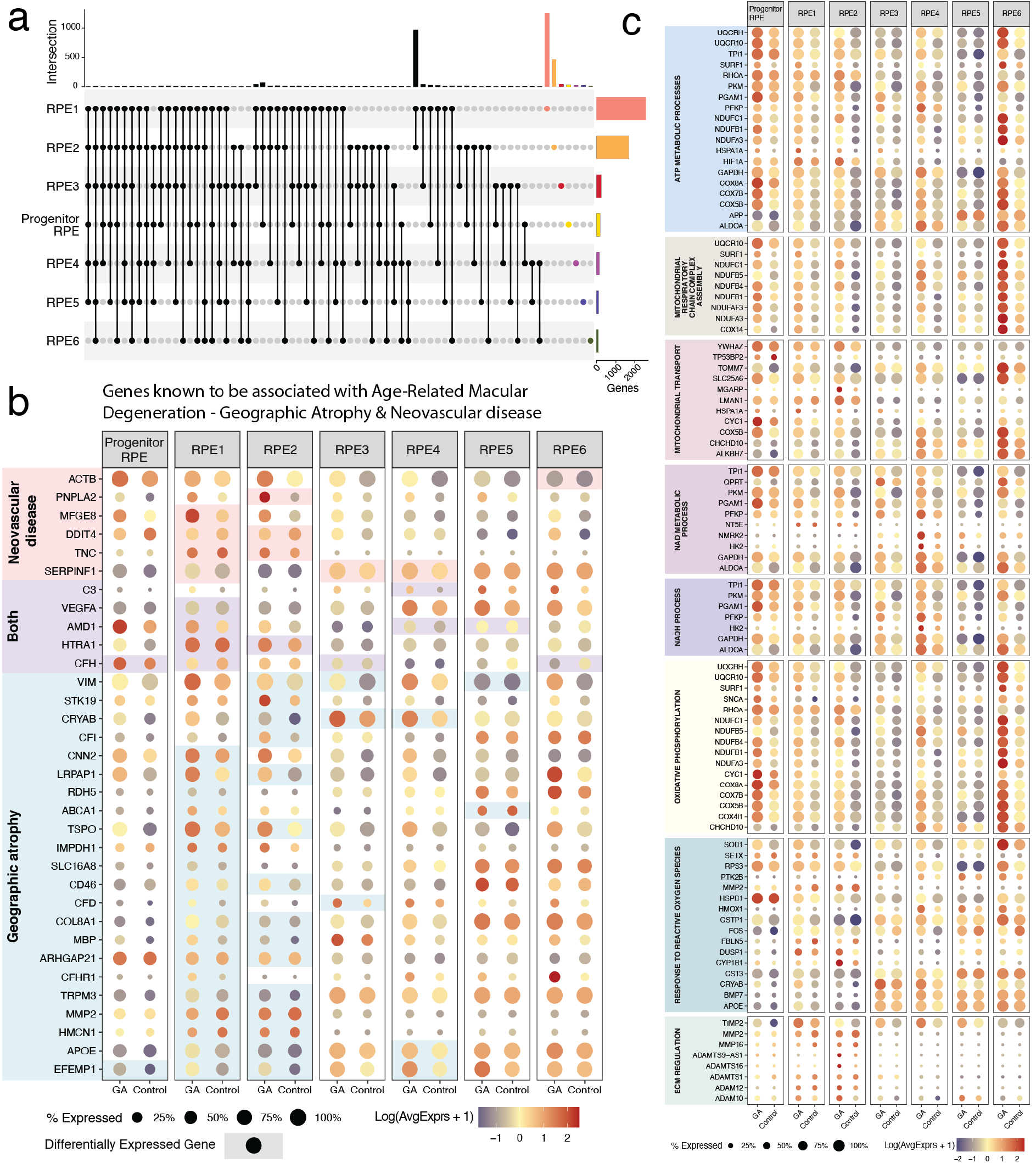
Genes associated with cell subpopulations and their expression in geographic atrophy and control cells. **(a)** Summary of cell subpopulation-specific gene regulation. **(b, c)** Dotplot representation of single cell expression profile of genes associated with geographic atrophy or neovascular AMD **(b)** and of genes associated with biological processes of interest **(c)** in geographic atrophy (GA) and in Control iPSC-derived subpopulations. Levels of gene expression per cell are shown with colour gradients, and frequencies of cells expressing the respective gene (% expressed) are shown with size of dots.

We performed Gene Ontology (GO)-ORA with disease-associated gene markers to identify biological processes, cellular components, and molecular functions that may be involved in the pathogenesis of AMD-GA. The geographic atrophy-progenitor RPE subpopulation was characterised by differential expression of genes involved in transcription, translation, and differentiation, including many ribosomal genes (**Supplementary Data 3**). The geographic atrophy-RPE1-5 subpopulations consistently showed differential expression of genes in various cellular component-, molecular- and biological process-pathways including in transcription and translation, protein localization to endoplasmic reticulum, ATP metabolic process, and apoptosis (**Supplementary Data 1**). RPE6 showed differential expression of genes mainly involved in transcription, translation and ribosome biogenesis as well as endoplasmic reticulum function (**Supplementary Data 1**). The amyloid fibril formation pathway was also differentially expressed in RPE2-4, whilst regulation of cell migration, and epithelial to mesenchymal transition (EMT) pathways were differentially expressed in RPE1-3,5 and RPE1,2,5, respectively (**Supplementary Data 1**). Genes associated with response to transforming growth factor beta and extracellular cellular matrix (ECM) reorganisation were also modified in the geographic atrophy RPE1 and RPE2 subpopulations (**Supplementary Data 3**). Interestingly, a substantial number of genes of the VEGFR signaling pathway was upregulated in RPE1 (*EMILIN1, NRP1, MYOF, ROCK2, HIF1A, PAK2, RHOA, CYBA, PIK3R1, PTK2B, CDC42, SHB, VEGFB, ITGA5, NCKAP1, BCAR1, NIBAN2, BAIAP2, CADM4, PTK2, VEGFA, ROCK1, VEGFC, SULF1, MAPKAPK2*, **Supplementary Data 3**). Of note, many genes coding for proteins involved in ECM regulation and known to play roles in retinal biology and in AMD ^44^ were differentially expressed in subpopulations of the geographic atrophy case cohort, including matrix metalloproteinases (MMPs), tissue inhibitors of metalloproteinases (TIMPs), a disintegrin and metalloproteinase domain (ADAMs), and a disintegrin and metalloproteinase with thrombospondin motifs (ADAMTSs). For instance, *TIMP2* was upregulated in geographic atrophy RPE1-3 whilst *TIMP3* was downregulated in RPE1 and upregulated in RPE2. Similarly, *MMP2* was upregulated in RPE1 and downregulated in RPE2 and *MMP16* was upregulated in both geographic atrophy subpopulations (Figure 3c, **Supplementary Data 3**). Mitochondrial activities such as oxidative phosphorylation, mitochondrial respiratory chain complex assembly and mitochondrial transport were increased in RPE1 and RPE2 (Figure 3c, **Supplementary Data 3**). Further, RPE1,2,5 were also characterized by modifications in genes involved in the ATP metabolic process, NAD metabolic process, and NADH process (Figure 3c, **Supplementary Data 3**). Finally, genes involved in the response to reactive oxygen species were upregulated in RPE1,3,4 (Figure 3c, **Supplementary Data 3**). At every step, our experimental workflow ensured that both control and geographic atrophy samples were assayed in shared conditions or randomized with respect to disease status (Methods). Hence, we are confident these transcriptomic differences are due to the genetic effects underlying the geographic atrophy risk. Moreover, environmental factors are unlikely to account for a difference in gene expression in differentiated cells, given the epigenetic profile of fibroblast-derived iPSCs is reset during reprogramming ^45, 46^.

### The proteomic analysis of RPE cells confirms specific protein expression in geographic atrophy cells

In parallel to the scRNA-Seq harvesting, all 79 lines were differentiated to RPE cells for mass spectrometry using a Tandem Mass Tag (TMT) platform for proteomics identification in control and geographic atrophy RPE cells (Figure 4, **Supplementary Data 1**). Given the experimental approach was not based on single cells but on a bulk harvest and analysis for each condition, cell cultures were assessed as a whole without distinction of subpopulations. Assessing protein expression levels, we observed that many of the proteins upregulated in the geographic atrophy cohort are typically involved in cell adhesion and ECM regulation (**Supplementary Data 1**). This observation was not obtained using pathway analysis at this stage. These include TIMP3 (the fifth-highest most upregulated protein in the geographic atrophy cohort), EFEMP1, ITGB4, SERPINB9; various tetraspanins (TSPAN6, TSPAN10, CD9/TSPAN29, CD82/TSPAN27), C1QTNF5 (second-highest most increased protein in the geographic atrophy cohort) and BSG. In the geographic atrophy RPE cells, the proteomic analysis also revealed increased levels of proteins known to be present in drusen ^47^, such as APOE and SCARB1 - also involved in cholesterol metabolism - and TIMP3. A number of signaling molecules were also upregulated in the geographic atrophy RPE cells, with increased levels of the growth factor SERPINF1/ PEDF (the fourth-highest upregulated protein in the geographic atrophy cohort); WNT signalling ligands SFRP1 and SFRP3/FRZB; the prostaglandin-synthesizing enzyme PTGDS, and the lysophosphatidic acid (LPA)-producing enzyme ENPP2/ATX. The complement pathway component C1R was also highly upregulated in the geographic atrophy RPE cells, as observed in other retinal dystrophies’ RPE cells ^48^. These data suggest that autocrine/paracrine signaling mechanisms are modified in the geographic atrophy RPE cells. No EMT markers were observed to be different between the two cohorts, suggesting that loss of epithelial cell features is not a hallmark of the geographic atrophy cohort cells that were examined. Proteins involved in various metabolic pathways were also upregulated in the geographic atrophy cohort, including for the retinoid metabolism (RETSAT, RDH11, RDH13), and reduced expression of the retinoic acid-binding protein CRABP1 - the most decreased protein in the geographic atrophy cohort), lipid metabolism (MLYCD, CYP20A1, CYP27A1, ACOT1, HSD17B12, TECR, KDSR, APOE, DHRS13), and gluconeogenesis and glycolysis (ALDOA, ENO3). Interestingly, the proteomic analysis revealed upregulation of many proteins from the respiratory chain pathway in geographic atrophy RPE. These include ATP5C1, SCO1 and mitochondrial complex I components (NDUFA3, NDUFA6, NDUFA8, NDUFA9, NDUFA10, NDUFA11, NDUFA13, NDUFB3, NDUFB5, NDUFB10, NDUFC2, NDUFS1, NDUFS3, NDUFV1, NDUFV2). Other enzymes involved in oxidoreductase activity were also upregulated, such as DHRS13, DHRS7B and GPX1. This may indicate that the geographic atrophy samples exhibit increased respiratory activity. Analysis of the dataset using STRING functional enrichment analysis (Biological Processes –Gene Ontology) supported this observation, confirming that specific pathways relating to mitochondrial processes were modulated in geographic atrophy cells. The dataset was highly enriched in pathways involved in oxidative phosphorylation, including mitochondrial electron transport (NADH to ubiquinone); mitochondrial respiratory chain complex I assembly; mitochondrial ATP synthesis coupled electron transport; oxidative phosphorylation; ATP metabolism; respiratory electron transport chain and reactive oxygen species (Figure 4a, **Supplementary Data 1**). Many of these same identified proteins were also represented in local network clustering (STRING) analysis (mitochondrial respiratory chain complex; oxidative phosphorylation and proton transporting) (Figure 4b, **Supplementary Data 1**). All markers associated with the pathways identified by GO and STRING analysis were upregulated in the geographic atrophy diseased cohort. Unsurprisingly, KEGG analysis found that oxidative phosphorylation was the most enriched biological process (ranked by strength parameters) in the diseased cohort, with many of the pathway hits closely related to the neurodegenerative diseases Parkinson’s and Alzheimer’s (**Supplementary Data 1**). Altogether, the large-scale proteomics analysis confirmed that proteins and pathways associated with ECM regulation, metabolism and mitochondrial functions are modified in geographic atrophy RPE cells.

**Figure 4.**
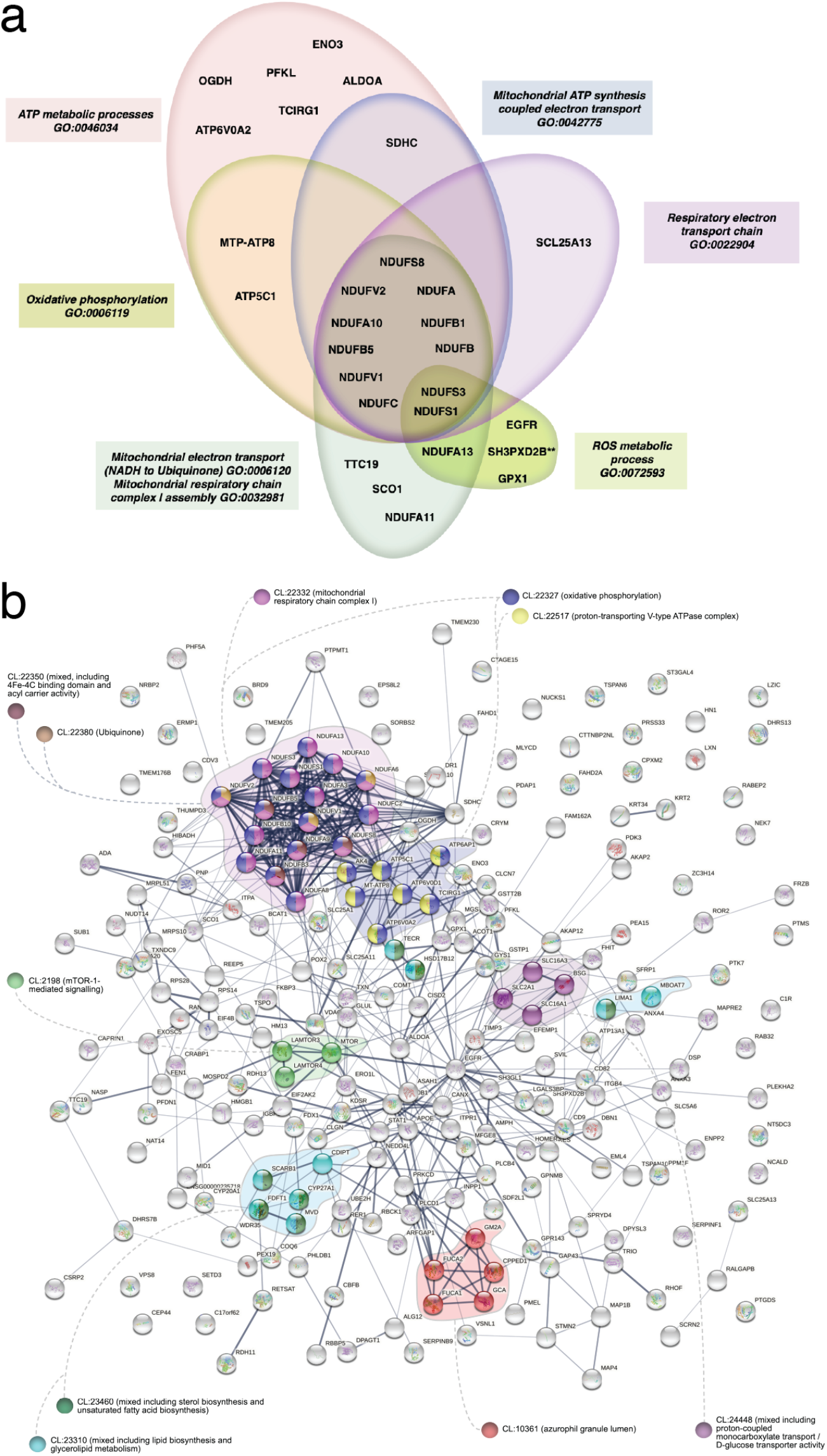
Proteome analysis of control and geographic atrophy-RPE cells. (**a**) Representation of enriched biological processes (Gene Ontology) related to mitochondrial function identified using STRING analysis. (**b**) Entire local network clustering (STRING) with enriched pathways highlighted in coloured nodules.

### RPE-associated genetic regulation of transcriptome and proteome

We investigated the relationship between genetic regulation and gene expression related to disease by mapping expression quantitative trait loci (eQTL) within each RPE subpopulation, and identified 439 cis-eQTL which surpassed the study-wide significance of FDR < 0.05 (Benjamini–Hochberg) and had a homozygous alternate allele in at least five individuals (Table 3, full results in **Table S6**). There is a high correlation between the detection of cis-eQTL and the number of cells in a subpopulation (Adjusted R^2^: 0.90, *p*-value: 6.3×10^-4^) which suggests the power to detect eQTL is related to the number of cells. The majority of eGenes (80.8%) - genes that had an eQTL - were subpopulation-exclusive, and only two eGenes - *GSTT1* and *RPS26 -* were common to all subpopulations (Figure 5a). The lead SNP for the eQTL at *GSTT1* was rs5760147 in most RPE subpopulations with the exception of Progenitor RPE (rs6003988) and RPE6 (rs2097433). *RPS26* was only associated with two variants - rs10876864 in RPE1, RPE5 and RPE6, and rs1131017 in Progenitor RPE, RPE2, RPE3 and RPE4. RPE1 and RPE2 share the greatest number of eQTLs (15) and eGenes (54) (**Table S7**). The effect sizes of these shared cis-eQTL are highly correlated (*r =* 0.99, *p*-value = 3.8×10^-14^), suggesting they are common genetic regulation mechanisms in these two subpopulations. We matched our results with previous studies and identified two eGenes with known associations with geographic atrophy - PRPH2 in RPE2 (rs9394903)^49^, and PILRB in RPE1 (rs11772580), RPE2 (rs11772580) and RPE3 (rs2404976)^27^ (Figure 5b). Lead eQTL identified in the preliminary round of mapping then underwent additional testing to identify interaction effects between alternative allelic effects and disease, and detected 45 significant interactions across all profiled subpopulations (p value: < 0.05) (**Table S6**).

**Table 3.** Summary of lead cis-eQTL per subpopulation.

| Subpopulation | Number of eQTLs | Number of eSNPs | Number of interactions |
| --- | --- | --- | --- |
| Progenitor RPE | 37 | 37 | 4 |
| RPE1 | 242 | 233 | 22 |
| RPE2 | 91 | 90 | 8 |
| RPE3 | 36 | 35 | 9 |
| RPE4 | 6 | 5 | 1 |
| RPE5 | 19 | 19 | 1 |
| RPE6 | 8 | 8 | 0 |
| Total | 439 | 377 | 45 |

**Figure 5.**
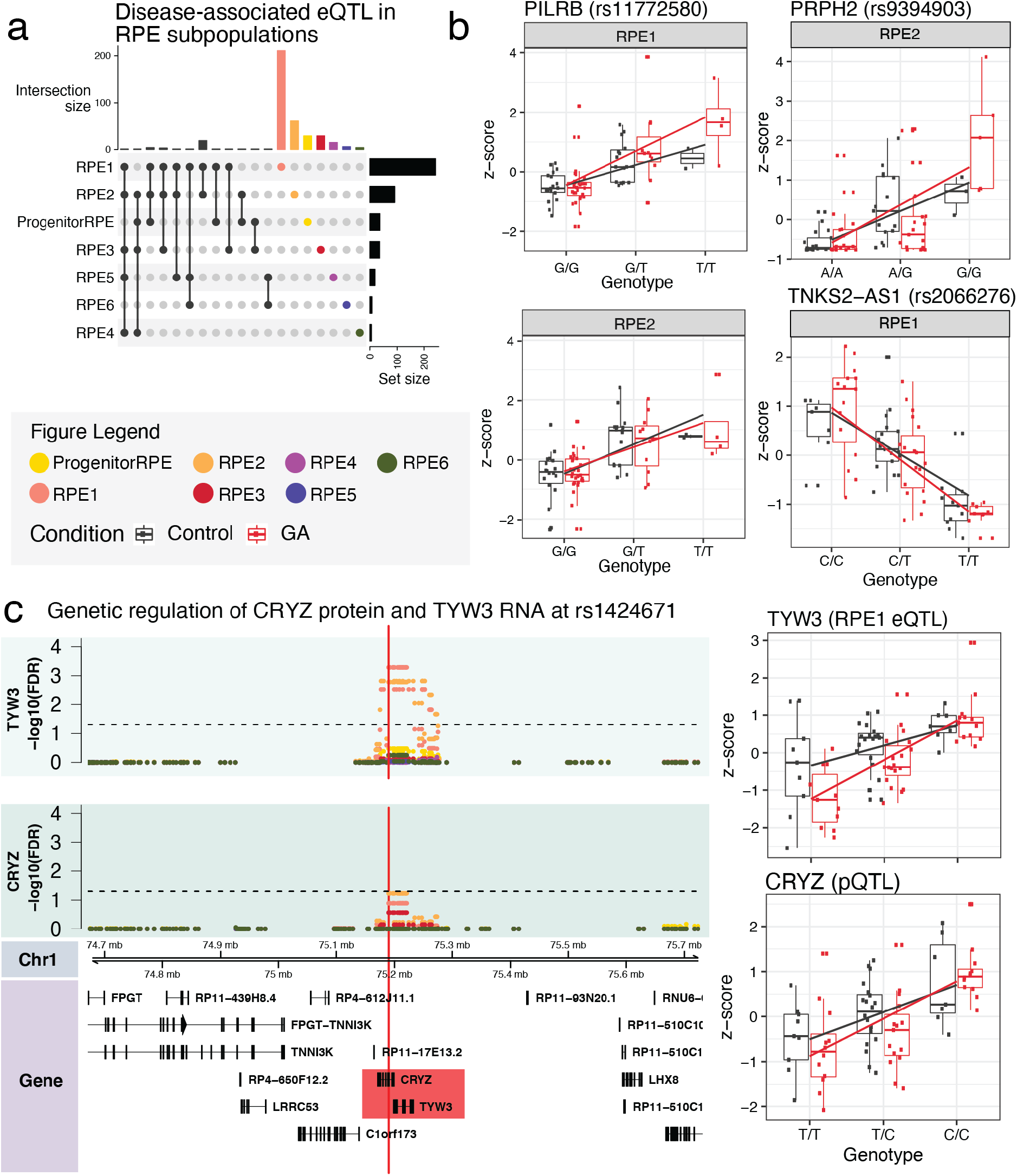
Genetic regulation of disease-associated expression and protein in RPE. **(a)** Distribution of disease-associated cis-eQTL across RPE subpopulations. **(b)** Gene-donor expression boxplots by genotype of eGenes that have a known association with AMD-GA - PILRB (RPE1 and RPE2), and PRPH2 (RPE2), and a candidate eGene TNKS2-AS1 (RPE1). (**c)** Locus zoom plot focusing on the 500kbp region around rs1424671 that includes TYW3 (eQTL interaction in RPE1) and CRYZ (pQTL interaction in bulk samples). Scatter plot axes show −log10(FDR) values of eQTL results of genes TYW3 and CRYZ across all RPE subpopulations in this region.

As protein expression does not necessarily correlate well with mRNA expression levels^50^, we performed cis-protein quantitative trait loci (cis-pQTL) to identify genetic variants that regulate protein expression in the RPE cells in the context of disease. It must, however, be noted that the proteomic analysis was performed on bulk RPE cultures, hence the identified pQTLs cannot be assigned to individual cells or subpopulations. We identified six proteins that share a lead SNP with RPE subpopulation-level eQTLs - PYROXD2/Q8N2H3, RNF13/O43567, CRYZ/Q08257, SPATA20/Q8TB22.2, PCOLCE/Q15113 and FIS1/Q9Y3D6 (Table 4). The variants associated with cis-pQTLs in PYROXD2, RNF13 and SPATA20 all occur with the same gene in corresponding cis-eQTLs, while the SNP associated with the cis-pQTL CRYZ - rs1424671, is associated with the eGene TYW3 from RPE1 and RPE2. Two pQTLs - PCOLCE and FIS1, have variants associated with the eGene CTA-339C12.1. SPATA20 (Spermatogenesis Associated 20)’s functions remain elusive. PYROXD2 (pyridine nucleotide-disulphide oxidoreductase domain 2) is a mitochondrial oxidoreductase regulating mitochondrial function and mitochondrial DNA copy number ^51^ and FIS1 (mitochondrial fusion protein 1) regulates mitochondrial dynamics, a process involved in various pathologies when dysregulated ^52^. RNF13 (Ring finger protein 13) is a crucial mediator of endoplasmic reticulum stress-induced apoptosis ^53^ and CRYZ (crystallin zeta, also known as quinone reductase) is an evolutionarily conserved protein induced under oxidative stress conditions ^54^ as well as detoxification of lipid peroxidation products ^55^, whilst PCOLCE (procollagen C-proteinase enhancer) is a collagen-binding protein involved in ECM formation and when dysregulated in fibrosis ^56^.

**Table 4.** Common genetic regulation mechanisms of transcriptome and proteome in RPE.

| Subpopulation (eQTL) | Uniprot ID | Gene (pQTL) | rsID | Beta (pQTL) | P-value (pQTL) | FDR (pQTL) | Gene (eQTL) | Beta (eQTL) | P-value (eQTL) | FDR (eQTL) |
| --- | --- | --- | --- | --- | --- | --- | --- | --- | --- | --- |
| RPE1 | Q08257 | CRYZ | rs1424671 | 5.97E-01 | 3.66E-03 | 2.56E-02 | TYW3 | 8.53E-01 | 2.77E-08 | 5.14E-04 |
| RPE2 | Q08257 | CRYZ | rs1424671 | 5.97E-01 | 3.66E-03 | 2.56E-02 | TYW3 | 8.60E-01 | 4.60E-08 | 1.65E-03 |
| RPE1 | Q8N2H3 | PYROXD2 | rs942813 | 6.37E-01 | 1.59E-03 | 1.67E-02 | PYROXD2 | 8.57E-01 | 1.60E-07 | 2.12E-03 |
| RPE1 | Q8TB22.2 | SPATA20 | rs989128 | -6.64E-01 | 1.59E-02 | 6.69E-02 | SPATA20 | -9.02E-01 | 1.10E-07 | 1.76E-04 |
| RPE1 | O43567 | RNF13 | rs772903 | -8.02E-01 | 1.67E-04 | 3.51E-03 | RNF13 | -7.40E-01 | 5.74E-06 | 4.54E-02 |
| RPE1 | Q15113 | PCOLCE | rs13239622 | -5.76E-01 | 1.37E-02 | 6.69E-02 | CTA-339C12.1 | 7.40E-01 | 5.86E-06 | 2.28E-02 |
| RPE1 | Q9Y3D6 | FIS1 | rs1859628 | 5.50E-01 | 2.60E-02 | 9.11E-02 | CTA-339C12.1 | 8.08E-01 | 2.87E-06 | 1.17E-02 |

### Transcriptome-wide association study analysis identifies novel genetic associations for geographic atrophy

Finally, we used the iPSC-derived RPE single cell eQTL data in conjunction with AMD GWAS summary statistics to prioritize AMD risk genes in a transcriptome-wide association study analysis (TWAS). In the single-cell TWAS, we identified 200 genes associated with AMD after Bonferroni correction in each RPE subpopulation, of which 38 were not genome-wide significant in the per-SNP analysis (best GWAS SNP P value < 5× 10^-8^) (Figure 6, **Supplementary Data 1**). Across different subpopulations, the TWAS results were generally consistent for several well-established regions, such as the *CFH* locus in chromosome 1, the *ARMS2/HTRA1* locus in chromosome 10 and *PILRB* in chromosome 7 (Figure 6, **Supplementary Data 1**). Interestingly, the most significantly associated transcript at the chromosome 10q26 locus varied from *HTRA1* in progenitor RPE and RPE6 cells to *ARMS2* in RPE2 cells. The *CFH* gene was significantly associated in all but the RPE3 and RPE4 subpopulations (Figure 6, **Supplementary Data 1**). Compared to a previous TWAS analysis based on bulk eQTL datasets ^28^, *PARP12* gene was also replicated in our single-cell TWAS in the RPE1 cell eQTL data (Figure 6, **Supplementary Data 1**). For the previously reported gene *RLBP1* ^28^, nearby gene *IDH2* was identified instead. Interestingly, the top GWAS SNP rs2238307 in gene *IDH2* is highly correlated with the top SNP rs3825991 in *PARP12* (r2 = 0.77) (**Supplementary Data 1**). rs10137367 at the *RDH11* locus was also identified in RPE1 by the single-cell TWAS (Figure 6, **Supplementary Data 1**).

**Figure 6.**
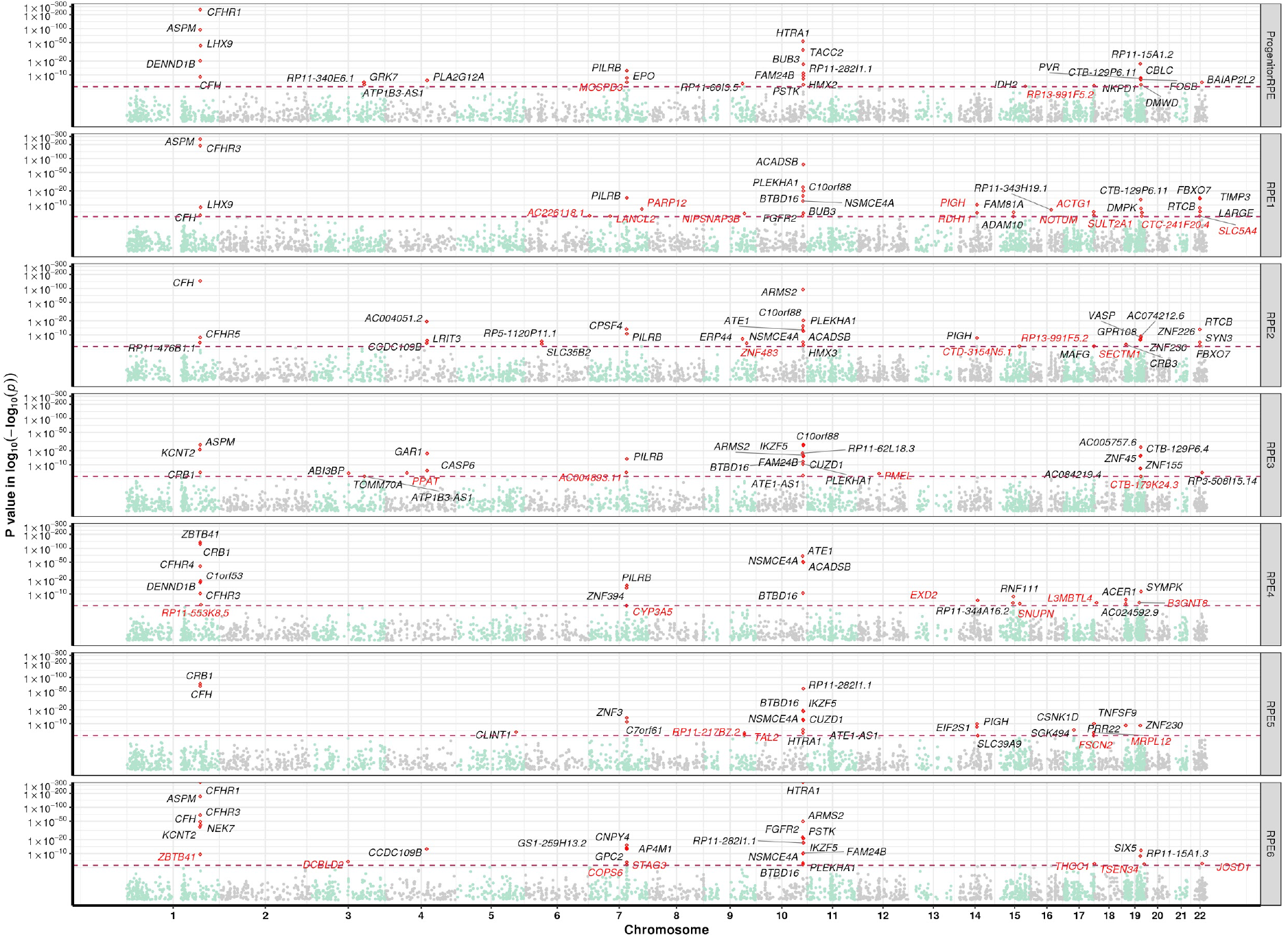
Prioritization of geographic atrophy risk genes. Genes that are significant after Bonferroni correction are highlighted with red dots, with the nearest gene names in black text (previously implicated genes), of which not genome-wide significant in the per-SNP analysis (top GWAS SNP *p*-value < 5× 10^-8^) are highlighted in red text (novel genes identified). The x-axis is the genome position from chromosome 1 to chromosome 22, the y-axis is the TWAS *p*-value in log-log scale. The maroon horizontal dash line is the Bonferroni correction level.

## Discussion

Here, we present a large-scale scRNA-seq analysis of iPSC-derived RPE cells affected in geographic atrophy. Following the capture of 204,126 cells, we analysed 127,659 cells from 79 individuals. The cell classification and trajectory analysis clearly indicated the efficiency of RPE differentiation, with most cells within the cultures identified as of RPE lineage. The variations observed within the RPE subpopulations were reflective to changes in maturity, rather than in cell identity, which is consistent with our previous work ^34^. Over 3,000 genes were differentially expressed between control and geographic atrophy RPE cells, with most of them in the two main RPE cell subpopulations RPE1 and RPE2. Genes with common risk alleles associated with geographic atrophy and neovascular AMD were upregulated in geographic atrophy RPE cells, including *CFH*, *HTRA1*, *EFEMP1 and APOE*, which provide further evidence of their involvement in the pathogenesis of geographic atrophy. Genes associated with specific biological pathways were upregulated in the geographic atrophy cohort, pointing to important functional differences between healthy and geographic atrophy RPE cells. In particular, our analysis revealed underlying differences in mitochondrial bioenergetic pathways, response to reactive oxygen species, ECM characteristics and autocrine/paracrine signaling. Key mitochondrial transcripts were altered in geographic atrophy RPE cells, with an increased expression of genes of the mitochondrial OXPHOS complex I machinery, oxidative phosphorylation, mitochondrial respiratory chain complex assembly and mitochondrial transport. Various metabolic pathways were also upregulated in the geographic atrophy cohort, in particular the ATP and the NAD/ NADH metabolic processes. The upregulation of genes involved in the response to reactive oxygen species in the geographic atrophy RPE cells further support a key role of cellular stress underlying RPE dysfunctions associated with geographic atrophy progression. The upregulation of genes involved in ECM reorganisation in the geographic atrophy samples corroborate current knowledge suggesting a role of the interaction of the RPE with the ECM in the development of geographic atrophy. Indeed, many of the genes we identified play roles in retinal biology and in AMD ^44^.

The scRNA-seq analysis is strongly supported by the large-scale proteomics analysis carried out using the same lines. Although performed with a bulk approach, which is unable to distinguish the various subpopulations of RPE identified by scRNA-Seq within samples, the analysis of the RPE proteome from both cohorts confirmed an upregulated expression of proteins involved in cell adhesion and ECM regulation. These included TIMP3, EFEMP1, ITGB4, SERPINB9, various tetraspanins, C1QTNF5 and BSG in the geographic atrophy RPE - many of which had already been identified by scRNA-Seq. Mutations in *TIMP3* are causative of the macular dystrophies Sorsby fundus dystrophy and in *EFEMP1 of* Doyne honeycomb retinal dystrophy and Malatia Leventiese, which are characterised by drusen accumulation underneath the RPE, an aspect that has been recapitulated *in vitro* using patient iPSC-derived RPE cells ^48^. Indeed, proteomic studies on drusen composition have identified TIMP3 and APOE as common constituents ^57^, both of which were upregulated in the geographic atrophy cohort. Tetraspanins are transmembrane molecular scaffolds that concentrate proteins into tetraspanin-enriched microdomains. By their interaction with molecules involved in adhesion, ECM regulation and cytoskeletal rearrangements, they play roles in various cellular processes, including adhesion, migration, and signalling ^58^ and have been implicated in various pathological events, such as angiogenesis in AMD, wound healing and immune cell response and inflammation ^59^. Variants in *C1QTNF5*, a membrane protein involved in cell adhesion and secretion, are associated with late-onset retinal degeneration ^60–63^. The immunoglobulin BSG, an extracellular matrix metalloproteinase inducer which stimulates cells to produce MMPs, plays roles in immune responses and has been implicated in photoreceptor survival ^64, 65^.

Interestingly, retinoid and pigmentation-related proteins were amongst the most significantly differentially expressed in the proteomic analysis comparing the geographic atrophy to healthy samples. The retinoid binding protein CRABP1 is the most decreased protein in the geographic atrophy cohort, which complements data showing decreased measurements of CRABP1 in late stage AMD eyes ^66^. By contrast, retinol dehydrogenase proteins RDH11 and RDH13, were highly upregulated in the geographic atrophy cohort. RDH13 is expressed in the retina but has no known role in the classical visual cycle, however elevated levels in the retinae of *Nrl^-/-^* mice suggest it may be important for cone function ^67^. By contrast, RDH11 directly participates in the visual retinoid cycle through the oxidation of 11-*cis* retinol ^68^. Pigmentation-related proteins PMEL and GPR143 were in greater abundance in the geographic atrophy cohort, suggesting factors relating to melanosome formation and function could be involved in disease. Indeed, melanosome movement and localisation at the apical surface of the RPE is known to play an important role in the maintenance of photoreceptor integrity ^69^. Taken together, our data suggests that RPE cells from the geographic atrophy cohort show constitutive differences in visual cycle processes and melanosome function to healthy RPE cells. Whether these differences are simply associated or causative of phenotypes remains to be assessed.

Various signaling molecules were also upregulated in the geographic atrophy RPE cells including the LPA-producing enzyme ENPP2/ATX. We previously demonstrated that the adult human eye secretes LPA in various locations, which suggests a role of ATX/LPA in the normal physiology of the eye and potentially in disease ^70^. We also showed that functional ATX is mainly secreted apically of human PSC-derived RPE cells, and that LPA regulates RPE homeostasis and photoreceptor functions ^70^. Our findings, together with the long list of LPA’s pleiotropic effects in various cell types ^71^, warrants further investigation on the role of ATX/LPA axis in geographic atrophy.

The large-scale analysis of the proteome also identified an overrepresentation of metabolism-related pathways in the geographic atrophy proteome, including pathways related to sterol, fatty acid and glycerolipid biosynthesis and metabolism and oxidative phosphorylation. The retina is amongst the body’s most metabolically active tissues, and results of this study suggest perturbations to metabolic homeostasis is a feature of geographic atrophy, either as a consequence or cause of the disease process. Many proteins related to mitochondrial functions were upregulated within the geographic atrophy cohort, including pathways relating to respiratory activity, oxidative phosphorylation and oxidoreduction. Indeed, mitochondrial pathways have been hypothesised to play a role in AMD ^72^, and provide potential targets for therapy ^73^. Furthermore, the over-representation of mTOR pathways in the geographic atrophy samples is worth noting, as the mTOR pathway has been shown to directly control mitochondrial processes ^74^, and when overactive can inhibit autophagy ^75^. Another nod to metabolic perturbation in macular degeneration is the long-standing hypothesis that chronic inflammation associated with the disease potentially disturbs the metabolic processes that occur between the RPE and the photoreceptors, leading to increased subretinal lactate concentrations, glycolysis deficit and increased ROS production ^76^. The results of the proteomic screen further confirms the validity of the findings obtained by scRNA-Seq and strongly points to a central involvement of metabolic or mitochondrial dysfunction and oxidative stress as underlying molecular events in the progression of geographic atrophy.

Our study identified a total of 439 eQTL across all RPE subpopulations, with most eQTL being cell population specific and 15 shared by RPE1 and RPE2. Two eGenes, *RPS26* and *GSTT1*, were common to all RPE subpopulations. Interestingly, *RPS26*, which encodes for proteins forming the small subunits of ribosomes, was also found to be ubiquitous in iPSC-derived retinal organoids ^32^ and has been associated with Type 1 diabetes ^77^. *GSTT1* encodes for a glutathione S transferase which is protective against oxidative stress, and has also been associated with disease, including ophthalmic conditions such as glaucoma ^78^, diabetic retinopathy ^79^ and cataract ^80^. Implications of *GSTT1* variants in AMD remains controversial ^81, 82^. Of interest, we observed an association of the coding polymorphism in *GSDMD* in the RPE1 and RPE2 subpopulations. This is interesting as gasdermin is a potential target for inflammatory conditions ^83^, and could thus be considered a novel target for treatment of geographic atrophy, especially as gasdermin D is elevated in eyes from geographic atrophy patients, where it plays a key role in the NLRP3 inflammasome activation and subsequent RPE death ^84^. Other eGenes of potential interest include *PILRB*, a known genetic variant in an AMD locus, also identified as an eQTL in RPE1, RPE2 and RPE3 ^21^.

Of all identified eQTL, 45 had a significant interaction between the SNP allelic effect and geographic atrophy. Identification of a significant eQTL in the long non-protein coding RNA *TNKS2-AS1* could suggest an impact of this allelic effect on genome regulation, including of *TNKS2* - a telomere-related gene. This is of interest given telomere length was previously demonstrated to be significantly different in geographic atrophy patients’ leukocytes ^85^. The identification of significant eQTL in mitochondrial proteins, such as MTG2, further suggests that variations in mitochondrial activities in the RPE might be at play in the progression of geographic atrophy. *TTC39C* was also identified as a significant eQTL, which encodes for a protein involved in cilia functions and for which a variant (rs9966620) has previously been associated with diabetic maculopathy (Meng et al., 2019). Finally, identification of *CTSF* as a significant interaction between the SNP allelic effect and geographic atrophy is also noteworthy, given mutations in *CTSF* are causative of ceroid lipofuscinosis, a disease associated with abnormal lysosomal lipofuscin storage (OMIM: 603539).

The investigation of regulatory mechanisms of protein expression by pQTL analysis shed light on six pQTL variant-protein interactions. In particular, PYROXD2 and FIS1 regulate mitochondrial functions via oxidoreductase activity ^51^ or mitochondrial dynamics ^52^. Previous GWAS identified genetic variants of PYROXD2 associated with urine trimethylamine concentration and cardiovascular disease ^86, 87^, type 2 diabetes and obesity ^88^. Gain of function variants for RNF13 have been associated with severe neurodegeneration leading to congenital microcephaly, epileptic encephalopathy, and cortical blindness ^89^. CRYZ is induced under oxidative stress ^54^ and detoxification of lipid peroxidation products ^55^, both molecular events already implicated in AMD pathogenesis ^90–92^. Recent GWAS have identified *CRYZ* as a susceptibility gene for insulin resistance ^93^ and amyotrophic lateral sclerosis ^94^. Lastly, PCOLCE is known for its involvement in fibrosis in various organs and a potential therapeutic target for this disease phenotype ^56^. These variants regulate protein expression and abundance in the RPE cells and thus further highlight the important role of genetic effects on protein expression in geographic atrophy.

Finally, the single-cell TWAS identified 200 genes associated with AMD, confirming known associations for AMD, including in the *CFH* and *ARMS2/HTRA1* loci, and also identifying 38 novel genes associated with geographic atrophy. One of these genes, *IDH2*, was identified as a novel association. *IDH2* is a key factor involved in reductive carboxylation of α-ketoglutarate in the RPE, and overexpression of IDH2 can protect against oxidative damage, supporting *IDH2* is a putative causal gene for AMD risk and single-cell TWAS is a potential effective approach for fine-mapping and identify drug targets ^95^. *RDH11* was also identified by TWAS, with its protein highly upregulated in the geographic atrophy cells as well. This is another interesting new candidate given some *RDH11* variants are associated with retinal dystrophies (OMIM: 607849).

In summary, we have identified important constitutive differences in RPE homeostasis associated with geographic atrophy when compared to healthy RPE cells. Outside the scope of this work but of clear importance, is the functional validation of the identified molecular targets and their ability to prevent or alter the course of geographic atrophy pathogenesis. Although other work recently reported on the transcriptomic and proteomic profiles of 151 independent iPSCs ^96^, this work is the first description of a population-scale analysis of the transcriptome and proteome of iPSC-derived RPE cells, as well as associated with geographic atrophy.

## Acknowledgments and Funding

We thank all participants who donated skin biopsies. This research was supported by National Health and Medical Research Council (NHMRC) Practitioner Fellowship (AWH), Senior Research Fellowship (AP, 1154389; SM, 1154543), Career Development Fellowship (JEP) and Investigator grant (JEP), by research grants from the Macular Disease Foundation Australia (RHG, AWH, JEP, AP), the NHMRC (research grant 1059369 RHG, AP, synergy grant 1181010, RHG, AP), the DHB Foundation (GEL, AP), retina Australia (AWH, AP), the Ophthalmic Research Institute of Australia (GEL, AWH, AP), NIH/NIGMS grant R01 GM132129 (JAP), the University of Melbourne and Operational Infrastructure Support from the Victorian Government.

## Author contributions

Conceptualization, A.S., M.D., G.E.L., J.P., A.W.H., A.P.; Methodology, A.S., M.D., G.E.L., J.E.P, A.W.H, A.P..; Investigation, A.S., M.D., G.E.L., H.H.L., D.H., M.M., X.H., D.N., L.R., L.G., J.A.P., V.G., C.L.C, U.N, A.M.S., R.Z., N.F., V.K.G.; Resources, L.C., L.S.K., D.A.M., G.B., N.V., R.H.G., A.W.H.; Data analysis, A.S., M.D., G.E.L., M.M., R.Z., X.H., S.M.G., J.E.P, A.W.H., A.P.; Writing - original draft, A.S., M.D., G.E.L., J.E.P, A.W.H., A.P.; Writing - review & editing, all authors.; Supervision and project administration, J.E.P, A.W.H., A.P.; Funding Acquisition, G.E.L., R.H.G, J.E.P., A.W.H, A.P.

## Online methods

### Participant recruitment

All participants gave informed written consent. This study was approved by the Human Research Ethics committees of the Royal Victorian Eye and Ear Hospital (11/1031H, 13/1151H-004), University of Melbourne (1545394), University of Tasmania (H0014124) UWA (EPS) as per the requirements of the NHMRC, in accordance with the Declarations of Helsinki. Cases who had advanced AMD with geographic atrophy in at least one eye and an age at first diagnosis over 50 years, were recruited through local ophthalmic clinics (mean ± SD age at recruitment: 83.8 ± 8.2 years). The control cohort has previously been described in ^32^, and had no manifest ophthalmic disease or drusen. The mean ± SD age at recruitment for participants was 69.8 ± 9.5 years. To ensure a diagnosis of AMD and not a monogenic retinal disease causing atrophy, all case participants had drusen identified on clinical examination. Dominantly inherited drusen phenotypes such as Sorsby fundus dystrophy, Doyne’s honeycomb dystrophy and Malattia Leventinese as well as fleck dystrophies such as Stargardt’s disease were excluded.

### Fibroblast culture

Punch biopsies were obtained from subjects over the age of 18 years. Fibroblasts were expanded, cultured and banked in DMEM with high glucose, 10% fetal bovine serum (FBS), L-glutamine, 100 U/mL penicillin and 100 μg/mL streptomycin (all from Thermo Fisher Scientific, USA). All cell lines were mycoplasma-free (MycoAlert mycoplasma detection kit, Lonza, Switzerland). Fibroblasts at passage (p) 2 were used for reprogramming.

### Generation, selection and iPSC maintenance

The maintenance and passaging of iPSCs were performed using a TECAN liquid handling platform as described ^97^. Briefly, iPSCs were generated by the nucleofection (Lonza) of episomal vectors expressing *OCT-4*, *SOX2*, *KLF4*, *L-MYC, LIN28* and shRNA against *p53* ^98^ in feeder- and serum-free conditions using TeSR™-E7™medium (Stem Cell Technologies) as described ^97^. Pluripotent cells were selected using anti-human TRA-1-60 Microbeads (Miltenyi) ^31^ and subsequently maintained onto vitronectin XF™-coated plates (Stem Cell Technologies) in StemFlex™ (Thermo Fisher Scientific), with media changes every second day and weekly passaging using ReLeSR™ (Stem Cell Technologies). Pluripotency was assessed by expression of the markers OCT3/4 (sc-5279, Santa Cruz Biotechnology) and TRA-1-60 (MA1-023-PE, Thermo Fisher Scientific; ab16288, Abcam) by immunocytochemistry, and virtual karyotyping by CNV array on all lines, as described in ^97^. Only geographic atrophy lines were generated for this study, as all control lines were already generated, and characterised in ^32^.

### Differentiation of iPSCs into RPE cells

RPE cells were generated as we described previously ^34^. Briefly, iPSCs were differentiated in E6 medium (Stem Cell Technologies) containing N2 supplement (Life Technologies), penicillin - streptomycin (Life Technologies) for 21-38 days (to reach RPE differentiation), switched to RPE medium (!MEM, 5% FBS, non-essential amino acids, N1 supplement, penicillin - streptomycin - glutamate, taurine - hydrocortisone - triiodothyronine) and cultured for a further 22-29 days with media changes every 2-3 days. Cells were then passaged with 0.25% trypsin-EDTA and plated onto growth factor-reduced Matrigel-coated plates (Corning) to enrich in RPE cells for an additional 30 days (76-88 days total).

### RPE cell harvest and single-cell preparation

RPE cells were harvested with 0.25% Trypsin-EDTA for 8 min, inactivated with RPE medium, and dissociated using manual trituration to yield a single-cell suspension. The cell suspension was centrifuged (5 minutes, 300 g, 4 °C), following which cells were resuspended in 1 mL of 0.1% BSA in PBS solution. Subsequently, cells were counted and assessed for viability with Trypan Blue, then pooled (eight samples maximum) at a concentration of 1000 live cells/μl (1*10^6 cells/mL).

### Single cell 3’ RNA-sequencing and pre-processing of transcriptome data

Multi-donor single-cell suspensions were prepared for scRNA-seq using the Chromium Single Cell 3′ Library & Gel bead kit (10x Genomics; PN-120237). Each pool was loaded onto individual wells of 10x Genomics Single Cell A Chip along with the reverse transcription (RT) master mix to generate single-cell gel beads in emulsion (GEMs). Reverse transcription was performed using a C1000 Touch Thermal Cycler with a Deep Well Reaction Module (Bio-Rad) as follows: 45 min at 53 °C; 5 min at 85 °C; hold 4 °C. cDNA was recovered and purified with DynaBeads MyOne Silane Beads (Thermo Fisher Scientific; catalog no. 37002D). Subsequently, it was amplified as follows: for 3 min at 98°C; 12× (for 15 sec at 98 °C; for 20 sec at 67 °C; for 60 sec at 72°C); for 60 sec at 72 °C; hold 4 °C followed recommended cycle number based on targeted cell number. Amplified cDNA was purified with SPRIselect beads (Beckman Coulter; catalog no. B23318) and underwent quality control following manufacturer’s instructions. Sequencing libraries for each pool were labelled with unique sample indices (SI) and combined for sequencing across two 2 x 150 cycle flow cells on an Illumina NovaSeq 6000 (NovaSeq Control Software v1.6) using S4 Reagent kit (catalog no. 20039236). Raw base calls from the sequencer then underwent demultiplexing, quality control, mapping and quantification with the Cell Ranger Single Cell Software Suite 3.1.0 by 10x Genomics (https://www.10xgenomics.com/). The *count* pipeline was run on each pool, with the target cell number set to 20,000 and demultiplexed reads mapped to the *Homo sapiens* reference *hg19*/*GRCh37* from ENSEMBL (release 75). The resulting transcriptome data for each pool then underwent quality control using the *Seurat* R package ^99^. Cells were removed if they did not meet the upper and lower thresholds calculated from 3 Median Absolute Deviations (MAD) of total UMI counts and number of detected genes, and if transcripts from mitochondrial genes exceeded 25% of total transcripts. Raw UMI counts from remaining cells then underwent normalization and scaling using the SCTransform function as implemented in Seurat ^100^.

### SNP genotype analysis and imputation

DNA was extracted from cell pellets using QIAamp DNA Mini Kit (QIAGEN, 51306) following the manufacturer’s instructions. DNA concentration was determined with a SimpliNano spectrophotometer (GE Life Sciences), diluted to a final concentration of 10-15 ng/µl and samples were genotyped on the UK Biobank Axiom™ Arrays at the Ramaciotti Centre for Genomics, Sydney, Australia. Genotype data were exported into PLINK data format using GenomeStudio PLINK Input Report Plug-in v2.1.4 and screened for SNP and individual call rates (<0.97), HWE failure (*p*<10^-6^), and MAF (<0.01). Samples with excess autosomal heterozygosity or with sex-mismatch were excluded. In addition, a genetic relationship matrix from all the autosomal SNPs were generated using the GCTA tool and one of any pair of individuals with estimated relatedness larger than 0.125 were removed from the analysis. Individuals with non-European ancestry were excluded outside of an “acceptable” box of +/- 6SD from the European mean in PC1 and PC2 in a SMARTPCA analysis. The 1000G Phase 3 population was used to define the axes, and the samples were projected onto those axes. Imputation was performed on each autosomal chromosome using the Michigan Imputation Server with the Haplotype Reference Consortium panel (HRC r1.1 2016) and run using Minimac3 and Eagle v2.3 ^101, 102^. Only SNPs with INFO > 0.8 and MAF > 0.1 were retained for downstream analyses.

### Demultiplexing of cell pools into individual donors

We used two SNP-based demultiplexing methods (demuxlet v0.1-beta ^103^ and souporcell v2.0 ^104^) and two transcription-based doublet detection methods (scrublet v0.2.1 ^105^ and DobuletDetection v2.5.2 ^106^) to identify droplets that contained one cell (singlets) and assign those cells to the correct donor. Droplets were considered singlets if they were classified as a singlet by all four softwares and were assigned to the same individual by both demuxlet and souporcell. For demuxlet, allele frequencies were first calculated with *popscle dsc-pileup* using all default parameters and known SNP locations based on imputed SNP genotypes overlapping exons. To classify doublets and assign singlets to each individual we ran *popscle demuxlet* using those pileup counts with default parameters except --geno_error_offset set to 0.1 and -- geno_error_coeff set to 0. Souporcell was used to classify droplets as doublets or singlets and to assign the singlets to clusters with *souporcell_pipeline.py* using default parameters and the --common_variants parameter to use just the known SNP locations overlapping exons that had been imputed for the individuals in the study. The SNP genotypes of the identified clusters were then correlated with the reference SNP genotypes. A cluster was assigned to a given individual if the correlation between them was the highest for both that individual and that cluster. Scrublet was run four different times using four different minimum variable percentile gene levels: 80, 85, 90 and 95 with all other default recommendations. The best variable percentile gene level was selected based on the best bimodal distribution of the simulated doublets with a reasonable doublet threshold (i.e. at the lowest point between the bimodal distribution). DoubletDetection detected doublets based on simulated transcriptional profiles by using the *doubletdetection.BoostClassifier* function with 50 iterations, *use_phenograph* set to False and *standard_scaling* set to True. The number of doublets per iteration were visualised to ensure convergence.

### Classification of cell subpopulations

Cells were assigned to RPE subpopulations using a supervised cell classification method called *scPred* v0.9^35^. A classifier was prepared from a reference dataset that had been characterised in a previous study^34^, and consisted of iPSC-derived RPE cells that had been profiled at two time points: 1 month and 12 months. Expression data from the reference was normalised using the *SCTransform* method from Seurat v3^100^, log2-transformed and scaled. The normalised values were then reduced to 15 Principal Components (PCs) with the *eigenDecompose* function from *scPred* and used to train the model with default settings. Classification was performed on each batch, and each cell was assigned a probability of belonging to a reference cluster based on the fit of its expression profile to the reference. To account for the transitional nature of cells from this experiment, we applied an adaptive threshold based on a cell having a prediction probability that lies within 0.3 standard deviations of the mean of all prediction probabilities of a reference subpopulation.

### Integration and dimensionality reduction of transcriptome data from multiple pools

Transcriptome data from all pools were combined and batch-normalized using integration methods from Seurat v3^99^. Integration anchors were selected from 2000 of the most variable genes using Canonical Correlation Analysis (CCA). As the individual datasets had been normalized with SCtransform on a cell-cell level, integration was performed with the argument ‘“normalization.method = “SCT”’. Dimensionality reduction using Principal Component Analysis (PCA) and Uniform Manifold Approximation Projection (UMAP) ^107^ was then performed using values produced by the integration process.

### Trajectory inference and pseudotime-based differential expression analysis

Trajectory inference - the identification of global lineage structures and the ordering of cellular states, was performed using *slingshot* v2.0.0^108^. Unnormalized, log-transformed UMI counts from 2000 of the most variable genes and the UMAP projection generated from the integration step were used by *slingshot* to build a minimum spanning tree across the subpopulations progressing from ProgenitorRPE, and calculate pseudotime across the trajectories. This information was used by *tradeSeq* v1.6.0^36^ to fit a generalized additive model to six knots, and apply the following differential expression tests: *associationTest* to identify trajectory- and condition-specific markers, *patternTest* and *diffEndTest* to identify genes with different transcription dynamics during the progression of the trajectory and at the end of the trajectory, and *conditionTest* to identify genes specific to disease status. Pseudotime-based differential expression results were corrected for multiple testing using Bonferroni correction, and the threshold for significance was 2.5×10^-5^.

### Differential expression analysis

Gene markers specific to disease status and subpopulation were identified using differential expression analysis, as implemented in MAST v1.16^109^. MAST was run through Seurat’s FindMarkers function on log-transformed, unnormalized UMI counts with the following latent variables: total UMI count (library size), pool number, age and sex. Bonferroni correction was used to correct p-values for multiple testing, and the threshold for significance was |Average log2 fold change| > 0.25 and adjusted p-value < 1.8×10^-5^.

### Gene Ontology and Disease Ontology Over-Representation Analysis

Gene sets were prepared for each subpopulation, from genes identified via pseudotime-based differential expression analysis and standard differential expression analysis. If fold change information was available, genes were also grouped by direction of regulation. Over-representation analysis was performed with the Gene Ontology^110, 111^ and Disease Ontology^112^ databases, as accessed through *clusterProfiler*^113^. P-values were corrected for multiple testing using Benjamini & Hochberg FDR and filtered for significance using a q-value threshold of 0.2.

### Transcriptome wide association study

TWAS was implemented in the FUSION pipeline (available at https://github.com/gusevlab/fusion_twas) ^114^. We first computed the single-cell eQTL weights using the “blup”, “lasso”, and “enet” models ^114^ in each subpopulation. Then, the single-cell gene-expressions were used in a TWAS using AMD GWAS summary statistics ^23^ to evaluate the association between inferred gene-expression and AMD. The TWAS *p*-values were adjusted for multiple testing using the Bonferroni method (approximately 19000 genes in each subpopulation).

### Preparation of protein samples

RPE cell cultures were lysed in RIPA buffer supplemented with phosphatase and protease inhibitors, sonicated with a probe sonicator (40 HZ × 2 pulses × 15 s), and insoluble debris were removed by centrifugation at 14,000 rpm for 15 min at 4°C, prior to measurement of protein contents by standard bicinchoninic acid assay (MicroBCA protein assay kit, Thermo Scientific). Solubilised proteins were reduced using 5 mM dithiothreitol and alkylated using 10 mM iodoacetamide. Proteins (150 µg) were initially digested at room temperature overnight using a 1:100 enzyme-to-protein ratio using Lys-C (Wako, Japan), followed by digestion with Trypsin (Promega, Madison, WI) for at least 4 hours at 37°C also at a 1:100 enzyme-to-protein ratio. Resultant peptides were acidified with 1% trifluoroacetic acid and purified using styrene divinylbenzene-reverse phase sulfonate (Empore) stage tips. The proteome was identified on a Tandem Mass Tag (TMT) platform (Progenetech, Sydney, Australia).

### TMT Labelling

8 independent 10 plex TMT experiments were carried out. Briefly, dried peptides from each sample were resuspended in 100 mM HEPES (pH 8.2) buffer and peptide concentration measured using the MicroBCA protein assay kit. Sixty micrograms of peptides from each sample was subjected to TMT labelling with 0.8 mg of reagent per tube. Labelling was carried out at room temperature for 1 h with continuous vortexing. To quench any remaining TMT reagent and reverse the tyrosine labelling, 8 μl of 5% hydroxylamine was added to each tube, followed by vortexing and incubation for 15 min at room temperature. For each of the respective ten plex experiments, the ten labelled samples were combined, and then dried down by vacuum centrifugation. Prior to High-pH reversed-phase fractionation, the digested and TMT-labelled peptide samples were cleaned using a reverse-phase C18 clean-up column (Sep-pak, Waters) and dried in vacuum centrifuge. The peptide mixture was resuspended in loading buffer (5 mM ammonia solution (pH 10.5), separated into a total of 96 fractions using an Agilent 1260 HPLC system equipped with quaternary pump, a degasser and a Multi-Wavelength Detector (MWD) (set at 210-, 214- and 280-nm wavelength). Peptides were separated on a 55-min linear gradient from 3 to 30% acetonitrile in 5 mM ammonia solution pH 10.5 at a flow rate of 0.3 mL/min on an Agilent 300 Extend C18 column (3.5 μm particles, 2.1 mm ID and 150 mm in length). The 96 fractions were finally consolidated into eight fractions. Each peptide fraction was dried by vacuum centrifugation, resuspended in 1% formic acid and desalted again using SDB-RPS (3M-Empore) stage tips.

### Liquid chromatography electrospray ionization tandem mass spectrometry (LC-ESI-MS/MS) data acquisition

Mass spectrometric data were collected on an Orbitrap Lumos mass spectrometer coupled to a Proxeon NanoLC-1200 UHPLC. The 100 µm capillary column was packed with 35 cm of Accucore 150 resin (2.6 μm, 150Å; ThermoFisher Scientific). The scan sequence began with an MS1 spectrum (Orbitrap analysis, resolution 60,000, 400−1600 Th, automatic gain control (AGC) target 4 x105, maximum injection time 50 ms). Data was acquired for 90 minutes per fraction. Analysis at the MS2 stage consisted of higher energy collision-induced dissociation (HCD), Orbitrap analysis with resolution of 50,000, automatic gain control (AGC) 1.25 x105, NCE (normalized collision energy) 37, maximum injection time 120 ms, and an isolation window at 0.5 Th. For data acquisition including FAIMS, the dispersion voltage (DV) was set at 5000V, the compensation voltages (CVs) were set at −40V, −60V, and −80V, and TopSpeed parameter was set at 1.5 sec per CV.

### Proteomic data analysis

Spectra were converted to mzXML via MSconvert. Database searching included all entries from the Human UniProt Database (downloaded: August 2019). The database was concatenated with one composed of all protein sequences for that database in the reversed order. Searches were performed using a 50-ppm precursor ion tolerance for total protein level profiling. The product ion tolerance was set to 0.2 Da. These wide mass tolerance windows were chosen to maximize sensitivity in conjunction with Comet searches and linear discriminant analysis. TMT tags on lysine residues and peptide N-termini (+229.163 Da for TMT) and carbamidomethylation of cysteine residues (+57.021 Da) were set as static modifications, while oxidation of methionine residues (+15.995 Da) was set as a variable modification. Peptide-spectrum matches (PSMs) were adjusted to a 1% false discovery rate (FDR). PSM filtering was performed using a linear discriminant analysis, as described previously and then assembled further to a final protein-level FDR of 1%. Proteins were quantified by summing reporter ion counts across all matching PSMs, also as described previously. Reporter ion intensities were adjusted to correct for the isotopic impurities of the different TMT reagents according to manufacturer specifications. The signal-to-noise (S/N) measurements of peptides assigned to each protein were summed and these values were normalized so that the sum of the signal for all proteins in each channel was equivalent to account for equal protein loading. Finally, each protein abundance measurement was scaled, such that the summed signal-to-noise for that protein across all channels equaled 100, thereby generating a relative abundance (RA) measurement. Investigation of protein–protein interactions and functional enrichment GO analysis of DE proteins were performed with STRING database version 11.0. STRING analysis was performed on the entire proteomics dataset, generating a network of interactions (based on both evidence of functional and physical interactions). Network lines represent the protein interaction score, which was set at a minimum medium confidence (0.4). Active interaction sources were based on text mining, experiments, databases, co-expression, neighbourhood, gene fusion and co-occurrence data.

### Mapping of expression and protein QTL

QTL mapping was performed using expression and protein measurements, and the genotype data of cell line donors that had been filtered for common SNPs (4,309,001 SNPs, Minor Allele Frequency > 10%). For eQTL, a donor-gene matrix was generated for each RPE subpopulation by taking the mean of normalized, corrected UMI counts for each gene from cells belonging to each donor that was present in the subpopulation. Genes that were expressed in less than 30% of the donor population were excluded. The resulting values were log-transformed and quantile normalised with the ‘normalizeBetweenArrays’ function from the limma R package ^115^. MatrixEQTL was run with an additive linear model using sex, age and the top six genotype principal components as covariates, and lead eQTL was selected based on the following thresholds: FDR (Benjamini–Hochberg procedure) < 0.05 and homozygous alternate allele frequency > 5. To identify eQTL that had alternative allelic effects under different disease statuses, we included an interaction term (SNP:disease status) in the original linear model for each eQTL identified by preliminary mapping and filtered by p-value < 0.05 of the interaction term. pQTL mapping was performed at a bulk level using protein abundance measurements taken from individual RPE cell lines. Abundance measurements were normalised using rank-based inverse normal transformation, and the protein-donor matrix was converted to an exon-donor matrix by matching protein identifiers and isoforms to exons listed in the Ensembl *Homo sapiens* Genes database ^116, 117^. SNPs that were within 1MB of exons were selected for mapping, which was performed with the *cis* function from QTLtools ^118^. As the same protein measurement was used for each exon belonging to a protein, the abundance measurements for each exon were grouped by protein and the mean value of each group - the measurement of each protein, was used for testing. pQTL results were matched to eQTL results by matching lead SNPs from pQTL analysis to SNPs with an eGene from cis-eQTL analysis. Benjamini & Hochberg FDR values for filtered pQTL results were calculated using adjusted beta p-values and filtered for significance using a threshold of 0.1.

## References

1. Chakravarthy, U. et al. Clinical risk factors for age-related macular degeneration: a systematic review and meta-analysis. BMC Ophthalmol. 10, 31 (2010).

2. Bird, A. C. et al. An international classification and grading system for age-related maculopathy and age-related macular degeneration. The International ARM Epidemiological Study Group. Surv. Ophthalmol. 39, 367–374 (1995).

3. Wong, W. L. et al. Global prevalence of age-related macular degeneration and disease burden projection for 2020 and 2040: a systematic review and meta-analysis. Lancet Glob Health 2, e106–16 (2014).

4. Rosenfeld, P. J. et al. Ranibizumab for neovascular age-related macular degeneration. N. Engl. J. Med. 355, 1419–1431 (2006).

5. Brown, D. M. et al. Ranibizumab versus verteporfin for neovascular age-related macular degeneration. N. Engl. J. Med. 355, 1432–1444 (2006).

6. Avery, R. L. et al. Intravitreal Bevacizumab (Avastin) for Neovascular Age-Related Macular Degeneration. Ophthalmology vol. 113 363–372.e5 (2006).

7. CATT Research Group et al. Ranibizumab and bevacizumab for neovascular age-related macular degeneration. N. Engl. J. Med. 364, 1897–1908 (2011).

8. Holz, F. G., Strauss, E. C., Schmitz-Valckenberg, S. & van Lookeren Campagne, M. Geographic atrophy: clinical features and potential therapeutic approaches. Ophthalmology 121, 1079–1091 (2014).

9. Sacconi, R., Corbelli, E., Querques, L., Bandello, F. & Querques, G. A Review of Current and Future Management of Geographic Atrophy. Ophthalmol Ther 6, 69–77 (2017).

10. Holz, F. G. et al. Efficacy and Safety of Lampalizumab for Geographic Atrophy Due to Age-Related Macular Degeneration: Chroma and Spectri Phase 3 Randomized Clinical Trials. JAMA Ophthalmol. 136, 666–677 (2018).

11. Yehoshua, Z. et al. Systemic complement inhibition with eculizumab for geographic atrophy in age-related macular degeneration: the COMPLETE study. Ophthalmology 121, 693–701 (2014).

12. Rosenfeld, P. J. et al. A Randomized Phase 2 Study of an Anti–Amyloid β Monoclonal Antibody in Geographic Atrophy Secondary to Age-Related Macular Degeneration. Ophthalmology Retina vol. 2 1028–1040 (2018).

13. Rosenfeld, P. J. et al. Emixustat Hydrochloride for Geographic Atrophy Secondary to Age-Related Macular Degeneration: A Randomized Clinical Trial. Ophthalmology 125, 1556–1567 (2018).

14. Schmitz-Valckenberg, S. et al. Natural History of Geographic Atrophy Progression Secondary to Age-Related Macular Degeneration (Geographic Atrophy Progression Study). Ophthalmology 123, 361–368 (2016).

15. Klein, R. J. Complement Factor H Polymorphism in Age-Related Macular Degeneration. Science vol. 308 385–389 (2005).

16. Edwards, A. O. Complement Factor H Polymorphism and Age-Related Macular Degeneration. Science vol. 308 421–424 (2005).

17. Haines, J. L. Complement Factor H Variant Increases the Risk of Age-Related Macular Degeneration. Science vol. 308 419–421 (2005).

18. Hageman, G. S. et al. From The Cover: A common haplotype in the complement regulatory gene factor H (HF1/CFH) predisposes individuals to age-related macular degeneration. Proceedings of the National Academy of Sciences vol. 102 7227–7232 (2005).

19. Jakobsdottir, J. et al. Susceptibility Genes for Age-Related Maculopathy on Chromosome 10q26. The American Journal of Human Genetics vol. 77 389–407 (2005).

20. Rivera, A. et al. Hypothetical LOC387715 is a second major susceptibility gene for age-related macular degeneration, contributing independently of complement factor H to disease risk. Hum. Mol. Genet. 14, 3227–3236 (2005).

21. Fritsche, L. G. et al. A large genome-wide association study of age-related macular degeneration highlights contributions of rare and common variants. Nat. Genet. 48, 134–143 (2016).

22. Yan, Q. et al. Genome-wide analysis of disease progression in age-related macular degeneration. Hum. Mol. Genet. 27, 929–940 (2018).

23. Han, X. et al. Genome-wide meta-analysis identifies novel loci associated with age-related macular degeneration. Journal of Human Genetics vol. 65 657–665 (2020).

24. Fritsche, L. G. et al. Seven new loci associated with age-related macular degeneration. Nat. Genet. 45, 433–9, 439e1–2 (2013).

25. Edwards, S. L., Beesley, J., French, J. D. & Dunning, A. M. Beyond GWASs: illuminating the dark road from association to function. Am. J. Hum. Genet. 93, 779–797 (2013).

26. Strunz, T. et al. A mega-analysis of expression quantitative trait loci in retinal tissue. PLoS Genet. 16, e1008934 (2020).

27. Orozco, L. D. et al. Integration of eQTL and a Single-Cell Atlas in the Human Eye Identifies Causal Genes for Age-Related Macular Degeneration. Cell Rep. 30, 1246–1259.e6 (2020).

28. Ratnapriya, R. et al. Author Correction: Retinal transcriptome and eQTL analyses identify genes associated with age-related macular degeneration. Nat. Genet. 51, 1067 (2019).

29. Takahashi, K. et al. Induction of pluripotent stem cells from adult human fibroblasts by defined factors. Cell 131, 861–872 (2007).

30. Yu, J. et al. Induced pluripotent stem cell lines derived from human somatic cells. Science 318, 1917–1920 (2007).

31. Crombie, D. E. et al. Development of a Modular Automated System for Maintenance and Differentiation of Adherent Human Pluripotent Stem Cells. SLAS Discov 22, 1016–1025 (2017).

32. Daniszewski, M. et al. Retinal ganglion cell-specific genetic regulation in primary open angle glaucoma. bioRxiv 2021.07.14.452417 (2021) doi:10.1101/2021.07.14.452417.

33. McCarthy, S. et al. A reference panel of 64,976 haplotypes for genotype imputation. Nat. Genet. 48, 1279–1283 (2016).

34. Lidgerwood, G. E. et al. Transcriptomic Profiling of Human Pluripotent Stem Cell-derived Retinal Pigment Epithelium over Time. Genomics Proteomics Bioinformatics (2020) doi:10.1016/j.gpb.2020.08.002.

35. Alquicira-Hernandez, J., Sathe, A., Ji, H. P., Nguyen, Q. & Powell, J. E. scPred: accurate supervised method for cell-type classification from single-cell RNA-seq data. Genome Biol. 20, 264 (2019).

36. Van den Berge, K., et al. Trajectory-based differential expression analysis for single-cell sequencing data. Nat. Commun. 11, 1201 (2020).

37. Liao, D. S. et al. Complement C3 Inhibitor Pegcetacoplan for Geographic Atrophy Secondary to Age-Related Macular Degeneration: A Randomized Phase 2 Trial. Ophthalmology 127, 186–195 (2020).

38. Sreekumar, P. G. et al. Intra-vitreal αB crystallin fused to elastin-like polypeptide provides neuroprotection in a mouse model of age-related macular degeneration. J. Control. Release 283, 94–104 (2018).

39. Krogh Nielsen, M., et al. Systemic Levels of Interleukin-6 Correlate With Progression Rate of Geographic Atrophy Secondary to Age-Related Macular Degeneration. Invest. Ophthalmol. Vis. Sci. 60, 202–208 (2019).

40. Haimovici, R. et al. Symptomatic abnormalities of dark adaptation in patients with EFEMP1 retinal dystrophy (Malattia Leventinese/Doyne honeycomb retinal dystrophy). Eye 16, 7–15 (2002).

41. Pilotto, E. et al. Müller cells and choriocapillaris in the pathogenesis of geographic atrophy secondary to age-related macular degeneration. Graefes Arch. Clin. Exp. Ophthalmol. 257, 1159–1167 (2019).

42. Piñero, J. et al. The DisGeNET knowledge platform for disease genomics: 2019 update. Nucleic Acids Res. 48, D845–D855 (2020).

43. Schriml, L. M. et al. Disease Ontology: a backbone for disease semantic integration. Nucleic Acids Res. 40, D940–6 (2012).

44. García-Onrubia, L. et al. Matrix Metalloproteinases in Age-Related Macular Degeneration (AMD). Int. J. Mol. Sci. 21, (2020).

45. Lister, R. et al. Hotspots of aberrant epigenomic reprogramming in human induced pluripotent stem cells. Nature 471, 68–73 (2011).

46. Polo, J. M. et al. A molecular roadmap of reprogramming somatic cells into iPS cells. Cell 151, 1617–1632 (2012).

47. Wang, L. et al. Abundant lipid and protein components of drusen. PLoS One 5, e10329 (2010).

48. Galloway, C. A. et al. Drusen in patient-derived hiPSC-RPE models of macular dystrophies. Proc. Natl. Acad. Sci. U. S. A. 114, E8214–E8223 (2017).

49. Smailhodzic, D. et al. Central areolar choroidal dystrophy (CACD) and age-related macular degeneration (AMD): differentiating characteristics in multimodal imaging. Invest. Ophthalmol. Vis. Sci. 52, 8908–8918 (2011).

50. Battle, A., et al. Genomic variation. Impact of regulatory variation from RNA to protein. Science 347, 664–667 (2015).

51. Wang, T. et al. Pyridine nucleotide-disulphide oxidoreductase domain 2 (PYROXD2): Role in mitochondrial function. Mitochondrion 47, 114–124 (2019).

52. Archer, S. L. Mitochondrial Dynamics — Mitochondrial Fission and Fusion in Human Diseases. New England Journal of Medicine vol. 369 2236–2251 (2013).

53. Arshad, M. et al. RNF13, a RING finger protein, mediates endoplasmic reticulum stress-induced apoptosis through the inositol-requiring enzyme (IRE1α)/c-Jun NH2-terminal kinase pathway. J. Biol. Chem. 288, 8726–8736 (2013).

54. Fernández, M. R. et al. Human and yeast zeta-crystallins bind AU-rich elements in RNA. Cell. Mol. Life Sci. 64, 1419–1427 (2007).

55. Porté, S. et al. Kinetic and structural evidence of the alkenal/one reductase specificity of human ζ-crystallin. Cell. Mol. Life Sci. 68, 1065–1077 (2011).

56. Lagoutte, P., Bettler, E., Vadon-Le Goff, S. & Moali, C. Procollagen C-proteinase enhancer-1 (PCPE-1), a potential biomarker and therapeutic target for fibrosis. Matrix Biology Plus 100062 (2021).

57. Crabb, J. W. et al. Drusen proteome analysis: an approach to the etiology of age-related macular degeneration. Proc. Natl. Acad. Sci. U. S. A. 99, 14682–14687 (2002).

58. Termini, C. M. & Gillette, J. M. Tetraspanins Function as Regulators of Cellular Signaling. Front. Cell Dev. Biol. 5, (2017).

59. Jiang, X., Zhang, J. & Huang, Y. Tetraspanins in Cell Migration. Cell Adh. Migr. 9, 406 (2015).

60. Stanton, C. M. et al. Novel pathogenic mutations in C1QTNF5 support a dominant negative disease mechanism in late-onset retinal degeneration. Sci. Rep. 7, 12147 (2017).

61. Shu, X. et al. Disease mechanisms in late-onset retinal macular degeneration associated with mutation in C1QTNF5. Human Molecular Genetics vol. 15 1680–1689 (2006).

62. Chekuri, A. et al. Late-onset retinal degeneration pathology due to mutations in CTRP5 is mediated through HTRA1. Aging Cell vol. 18 (2019).

63. Kellner, U. et al. Autosomal Dominant Gyrate Atrophy-Like Choroidal Dystrophy Revisited: 45 Years Follow-Up and Association with a Novel Missense Variant. Int. J. Mol. Sci. 22, (2021).

64. Aït-Ali, N. et al. Rod-Derived Cone Viability Factor Promotes Cone Survival by Stimulating Aerobic Glycolysis. Cell vol. 161 817–832 (2015).

65. Muramatsu, T. Basigin (CD147), a multifunctional transmembrane glycoprotein with various binding partners. J. Biochem. 159, 481–490 (2016).

66. Nordgaard, C. L. et al. Proteomics of the Retinal Pigment Epithelium Reveals Altered Protein Expression at Progressive Stages of Age-Related Macular Degeneration. Invest. Ophthalmol. Vis. Sci. 47, 815–822 (2006).

67. Kanan, Y. et al. Retinoid processing in cone and Müller cell lines. Exp. Eye Res. 86, 344–354 (2008).

68. Parker, R. O. & Crouch, R. K. Retinol dehydrogenases (RDHs) in the visual cycle. Exp. Eye Res. 91, 788–792 (2010).

69. Burgoyne, T., O’Connor, M. N., Seabra, M. C., Cutler, D. F. & Futter, C. E. Regulation of melanosome number, shape and movement in the zebrafish retinal pigment epithelium by OA1 and PMEL. J. Cell Sci. 128, 1400–1407 (2015).

70. Lidgerwood, G. E. et al. Role of lysophosphatidic acid in the retinal pigment epithelium and photoreceptors. Biochim. Biophys. Acta Mol. Cell Biol. Lipids 1863, 750–761 (2018).

71. Frisca Frisca, F., Sabbadini, R. A., Goldshmit, Y. & Pébay, A. Biological Effects of Lysophosphatidic Acid in the Nervous System. International Review of Cell and Molecular Biology Volume 296 273–322 (2012) doi:10.1016/b978-0-12-394307-1.00005-9.

72. Golestaneh, N. et al. Repressed SIRT1/PGC-1α pathway and mitochondrial disintegration in iPSC-derived RPE disease model of age-related macular degeneration. J. Transl. Med. 14, 344 (2016).

73. Terluk, M. R. et al. Investigating mitochondria as a target for treating age-related macular degeneration. J. Neurosci. 35, 7304–7311 (2015).

74. Ramanathan, A. & Schreiber, S. L. Direct control of mitochondrial function by mTOR. Proc. Natl. Acad. Sci. U. S. A. 106, 22229–22232 (2009).

75. Nazio, F. et al. mTOR inhibits autophagy by controlling ULK1 ubiquitylation, self-association and function through AMBRA1 and TRAF6. Nat. Cell Biol. 15, 406–416 (2013).

76. Léveillard, T., Philp, N. & Sennlaub, F. Is Retinal Metabolic Dysfunction at the Center of the Pathogenesis of Age-related Macular Degeneration? International Journal of Molecular Sciences vol. 20 762 (2019).

77. Dixon, A. L. et al. A genome-wide association study of global gene expression. Nat. Genet. 39, 1202–1207 (2007).

78. Yu, Y., Weng, Y., Guo, J., Chen, G. & Yao, K. Association of glutathione S transferases polymorphisms with glaucoma: a meta-analysis. PLoS One 8, e54037 (2013).

79. Sun, L., Zhang, Y. & Xiong, Y. GSTM1 and GSTT1 null genotype and diabetic retinopathy: a meta-analysis. Int. J. Clin. Exp. Med. 8, 1677–1683 (2015).

80. Sun, W., Su, L., Sheng, Y., Shen, Y. & Chen, G. Is there association between Glutathione S Transferases polymorphisms and cataract risk: a meta-analysis? BMC Ophthalmology vol. 15 (2015).

81. Liu, M. M., Chan, C.-C. & Tuo, J. Genetic mechanisms and age-related macular degeneration: common variants, rare variants, copy number variations, epigenetics, and mitochondrial genetics. Hum. Genomics 6, 13 (2012).

82. Hunter, A. A., 3rd et al. GSTM1 and GSTM5 Genetic Polymorphisms and Expression in Age-Related Macular Degeneration. Curr. Eye Res. 41, 410–416 (2016).

83. Minton, K. Inflammasome: Looking death in the eye. Nat. Rev. Immunol. 18, 4 (2017).

84. Kerur, N. et al. cGAS drives noncanonical-inflammasome activation in age-related macular degeneration. Nat. Med. 24, 50–61 (2017).

85. Banevicius, M. et al. Association of relative leukocyte telomere length and genetic variants in telomere-related genes () with atrophic age-related macular degeneration. Ophthalmic Genet. 42, 189–194 (2021).

86. Nicholson, G. et al. A genome-wide metabolic QTL analysis in Europeans implicates two loci shaped by recent positive selection. PLoS Genet. 7, e1002270 (2011).

87. Wang, Z. et al. Gut flora metabolism of phosphatidylcholine promotes cardiovascular disease. Nature 472, 57–63 (2011).

88. Fadason, T., Ekblad, C., Ingram, J. R., Schierding, W. S. & O’Sullivan, J. M. Physical Interactions and Expression Quantitative Traits Loci Identify Regulatory Connections for Obesity and Type 2 Diabetes Associated SNPs. Front. Genet. 8, 150 (2017).

89. Edvardson, S. et al. Heterozygous RNF13 Gain-of-Function Variants Are Associated with Congenital Microcephaly, Epileptic Encephalopathy, Blindness, and Failure to Thrive. Am. J. Hum. Genet. 104, 179–185 (2019).

90. Kopitz, J., Holz, F. G., Kaemmerer, E. & Schutt, F. Lipids and lipid peroxidation products in the pathogenesis of age-related macular degeneration. Biochimie 86, 825–831 (2004).

91. Kaemmerer, E., Schutt, F., Krohne, T. U., Holz, F. G. & Kopitz, J. Effects of lipid peroxidation-related protein modifications on RPE lysosomal functions and POS phagocytosis. Invest. Ophthalmol. Vis. Sci. 48, 1342–1347 (2007).

92. Zhao, T., Guo, X. & Sun, Y. Iron Accumulation and Lipid Peroxidation in the Aging Retina: Implication of Ferroptosis in Age-Related Macular Degeneration. Aging Dis. 12, 529–551 (2021).

93. Qi, Q. et al. Genome-wide association analysis identifies TYW3/CRYZ and NDST4 loci associated with circulating resistin levels. Hum. Mol. Genet. 21, 4774–4780 (2012).

94. Wei, L. et al. Identification of TYW3/CRYZ and FGD4 as susceptibility genes for amyotrophic lateral sclerosis. Neurology Genetics vol. 5 e375 (2019).

95. Du, J. et al. Reductive carboxylation is a major metabolic pathway in the retinal pigment epithelium. Proc. Natl. Acad. Sci. U. S. A. 113, 14710–14715 (2016).

96. Mirauta, B. A. et al. Population-scale proteome variation in human induced pluripotent stem cells. Elife 9, (2020).

97. Daniszewski, M. et al. Single-Cell Profiling Identifies Key Pathways Expressed by iPSCs Cultured in Different Commercial Media. iScience 7, 30–39 (2018).

98. Okita, K. et al. A more efficient method to generate integration-free human iPS cells. Nat. Methods 8, 409–412 (2011).

99. Stuart, T. et al. Comprehensive Integration of Single-Cell Data. Cell 177, 1888–1902.e21 (2019).

100. Hafemeister, C. & Satija, R. Normalization and variance stabilization of single-cell RNA-seq data using regularized negative binomial regression. Genome Biol. 20, 296 (2019).

101. Das, S. et al. Next-generation genotype imputation service and methods. Nat. Genet. 48, 1284–1287 (2016).

102. Loh, P.-R. et al. Reference-based phasing using the Haplotype Reference Consortium panel. Nat. Genet. 48, 1443–1448 (2016).

103. Kang, H. M. et al. Multiplexed droplet single-cell RNA-sequencing using natural genetic variation. Nat. Biotechnol. 36, 89–94 (2018).

104. Heaton, H. et al. Souporcell: robust clustering of single-cell RNA-seq data by genotype without reference genotypes. Nat. Methods 17, 615–620 (2020).

105. Wolock, S. L., Lopez, R. & Klein, A. M. Scrublet: Computational Identification of Cell Doublets in Single-Cell Transcriptomic Data. Cell Syst 8, 281–291.e9 (2019).

106. Shor, J. DoubletDetection. (Github).

107. Becht, E. et al. Dimensionality reduction for visualizing single-cell data using UMAP. Nat. Biotechnol. (2018) doi:10.1038/nbt.4314.

108. Street, K. et al. Slingshot: cell lineage and pseudotime inference for single-cell transcriptomics. BMC Genomics 19, 477 (2018).

109. Finak, G. et al. MAST: a flexible statistical framework for assessing transcriptional changes and characterizing heterogeneity in single-cell RNA sequencing data. Genome Biol. 16, 278 (2015).

110. Ashburner, M. et al. Gene ontology: tool for the unification of biology. The Gene Ontology Consortium. Nat. Genet. 25, 25–29 (2000).

111. Gene Ontology Consortium. The Gene Ontology resource: enriching a GOld mine. Nucleic Acids Res. 49, D325–D334 (2021).

112. Schriml, L. M., et al. Human Disease Ontology 2018 update: classification, content and workflow expansion. Nucleic Acids Res. 47, D955–D962 (2019).

113. Yu, G., Wang, L.-G., Han, Y. & He, Q.-Y. clusterProfiler: an R package for comparing biological themes among gene clusters. OMICS 16, 284–287 (2012).

114. Gusev, A. et al. Integrative approaches for large-scale transcriptome-wide association studies. Nature Genetics vol. 48 245–252 (2016).

115. Ritchie, M. E. et al. limma powers differential expression analyses for RNA-sequencing and microarray studies. Nucleic Acids Res. 43, e47 (2015).

116. Smedley, D. et al. BioMart – biological queries made easy. BMC Genomics 10, 1–12 (2009).

117. Hubbard, T. et al. The Ensembl genome database project. Nucleic Acids Res. 30, 38–41 (2002).

118. Delaneau, O. et al. A complete tool set for molecular QTL discovery and analysis. Nat. Commun. 8, 15452 (2017).

