## Supplemental Information for "Transcriptomic and proteomic retinal pigment epithelium signatures of age-related macular degeneration"

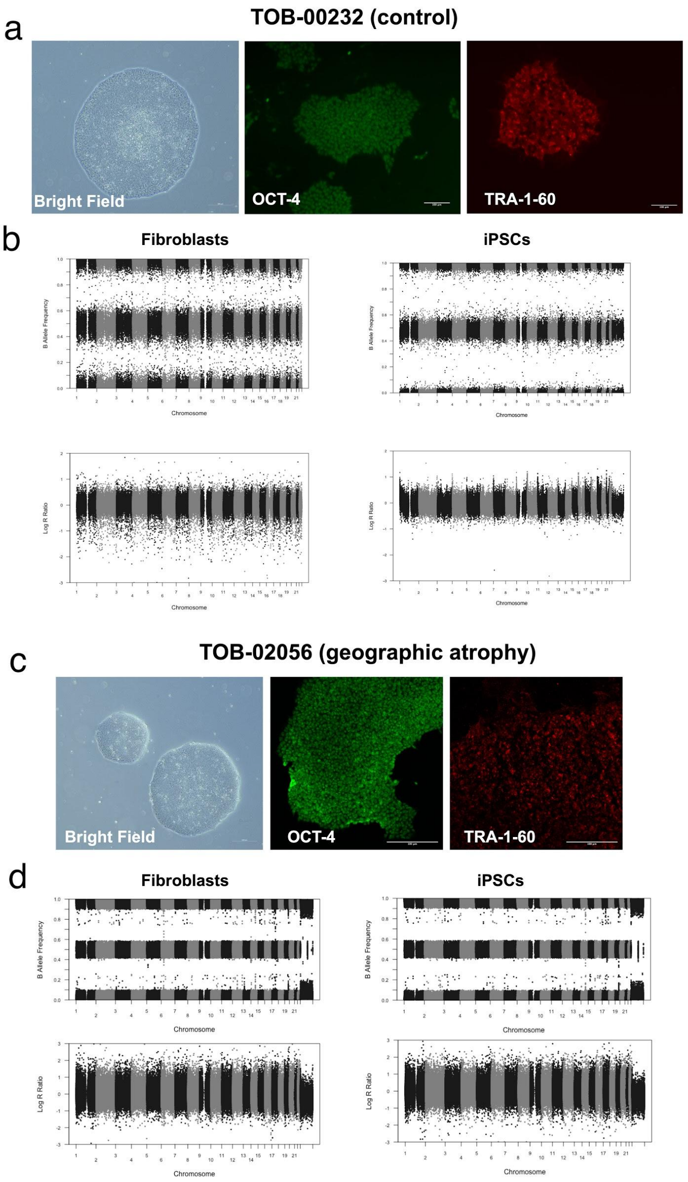


**Figure S1. Examples of quality control for iPSC lines, Related to Figure 1.** Control line TOB-00232 (**a, b**) and geographic atrophy line TOB-02056 (**c, d**) are shown. (**a, c**) Representative bright field images and immunostainings for the pluripotent markers OCT-4 and TRA-1-60 for both lines (Scale bars: 200 μm for bright field, and 100 μm for immunofluorescence). Images are representative of all cell lines. (**b, d**) Copy Number Variation Analysis for both parental fibroblasts and iPSCs representative of a normal virtual karyotyping. Each panel shows the B allele frequency (BAF) and the log R ratio (LRR). BAF at values other than 0, 0.5 or 1 indicate an abnormal copy number. LRR represents a logged ratio of “observed probe intensity to expected intensity”. A deviation from zero corresponds to a change in copy number.

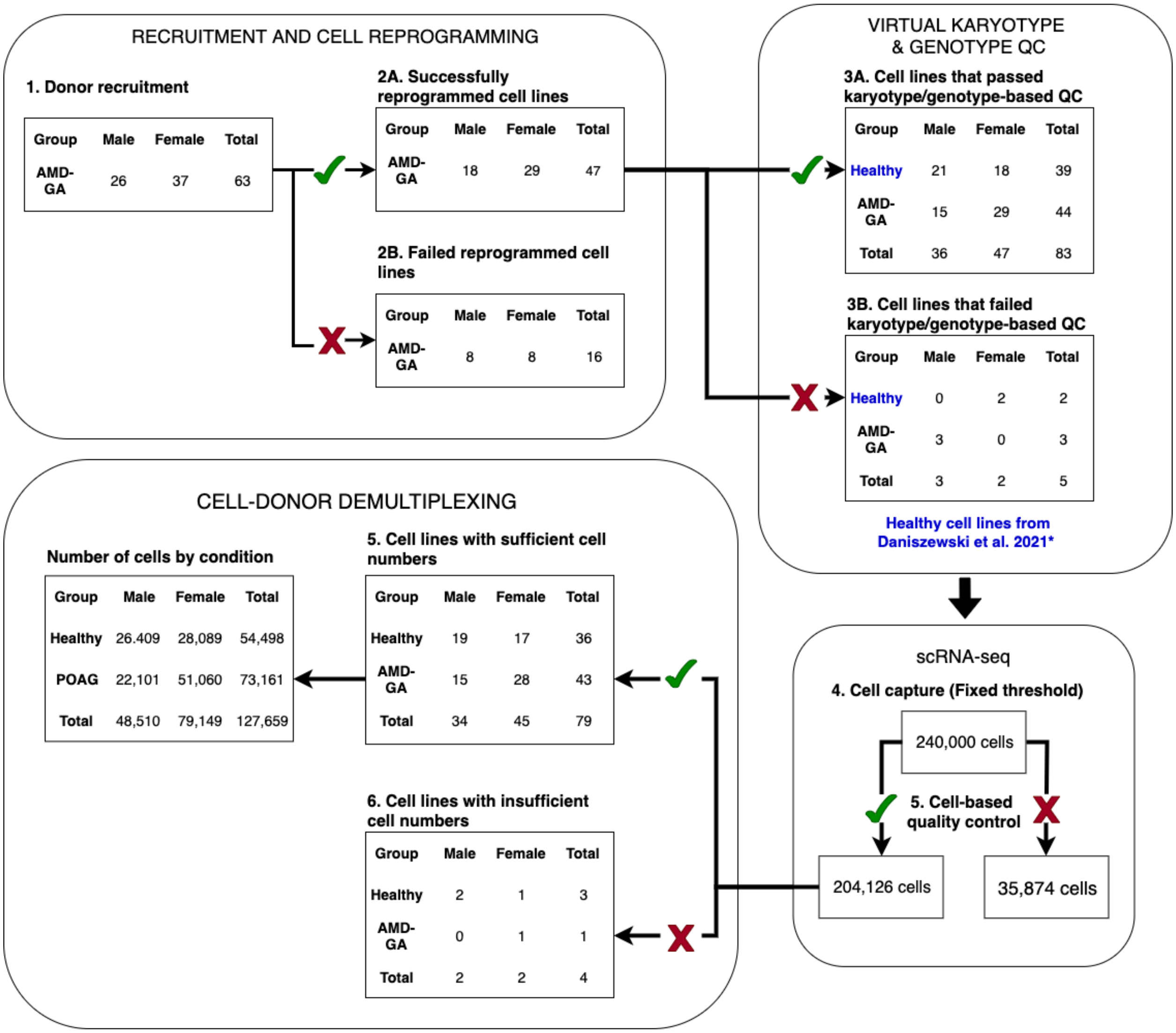


**Figure S2. Cell line and capture quality control flowchart.** Cell lines were generated from skin biopsies of AMD-GA patients, and reprogrammed into iPSCs and subsequently, RPE. Lines were discarded if they failed virtual karyotype and/or genotype QC, and if less than 200 cells were captured by scRNA-seq.

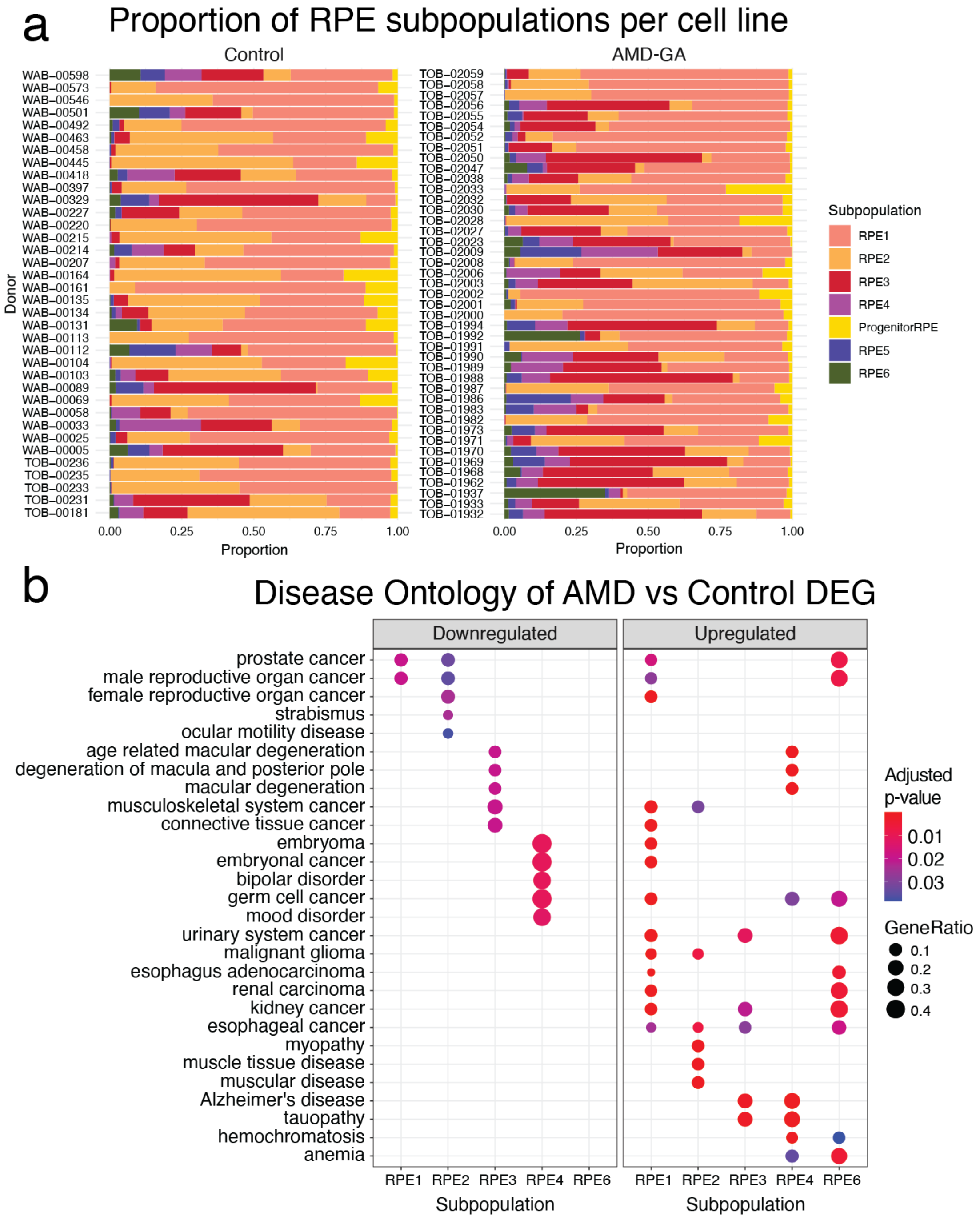


**Figure S3. Proportion of RPE subpopulations per cell line. (a)** Cells were matched to reference RPE subpopulations via cell type classification, and to donors via genotyping. Proportions of RPE subpopulations were calculated for each cell line. (**b)** Differentially expressed genes from AMD-GA cells of each RPE subpopulation underwent Disease Ontology (DO) and enrichment analysis. Upregulated and downregulated genes were analysed separately. GeneRatio represents proportion of genes in query dataset that were also present DO gene set. The colour scale represents the significance of the DO result.

**Table S1. Single cell RNA-sequencing quality metrics**

| Pool | 1 | 2 | 3 | 4 | 5 | 6 | 7 | 8 | 9 | 10 | 11 | 12 |
|---|---|---|---|---|---|---|---|---|---|---|---|---|
| Forced Number of Cells | 20,000 | 20,000 | 20,000 | 20,000 | 20,000 | 20,000 | 20,000 | 20,000 | 20,000 | 20,000 | 20,000 | 20,000 |
| Mean Reads per Cell | 22,427 | 22,144 | 21,305 | 22,104 | 22,387 | 47,134 | 45,139 | 45,698 | 45,068 | 44,226 | 45,159 | 46,388 |
| Median Genes per Cell | 2,516 | 2,426 | 2,519 | 2556 | 2,460 | 3,283 | 3,049 | 3,162 | 3,210 | 3,052 | 2,034 | 3,084 |
| Number of Reads | 448,558,250 | 442,888,016 | 426,111,365 | 442,084,959 | 447,748,740 | 942,691,103 | 902,785,800 | 913,970,169 | 901,377,134 | 884,523,529 | 903,190,965 | 927,769,625 |
| Valid Barcodes | 97.70% | 97.80% | 97.70% | 97.70% | 97.60% | 97.50% | 97.40% | 97.40% | 97.50% | 97.70% | 97.70% | 97.40% |
| Sequencing Saturation | 29.50% | 30.00% | 26.60% | 25.60% | 30.10% | 48.60% | 45.60% | 43.10% | 42.40% | 50.10% | 62.10% | 59.10% |
| Q30 Bases in Barcode | 96.30% | 96.30% | 96.30% | 96.30% | 96.40% | 95.80% | 95.80% | 95.80% | 95.80% | 95.80% | 95.80% | 95.80% |
| Q30 Bases in RNA Read | 94.80% | 94.70% | 95.00% | 94.90% | 95.00% | 94.40% | 94.10% | 94.40% | 94.20% | 94.00% | 94.20% | 94.40% |
| Q30 Bases in Sample Index | 93.50% | 88.40% | 94.60% | 95.30% | 90.30% | 94.20% | 93.40% | 89.80% | 89.20% | 94.20% | 92.10% | 94.80% |
| Q30 Bases in UMI | 96.20% | 96.20% | 96.10% | 96.10% | 96.20% | 95.60% | 95.50% | 95.60% | 95.60% | 95.60% | 95.50% | 95.50% |
| Reads Mapped to Genome | 96.60% | 97.00% | 97.00% | 96.80% | 97.10% | 97.30% | 96.90% | 96.90% | 97.00% | 97.20% | 97.30% | 97.60% |
| Reads Mapped Confidently to Genome | 93.80% | 94.30% | 93.50% | 93.40% | 94.50% | 94.80% | 94.30% | 94.30% | 94.60% | 95.00% | 94.80% | 95.20% |
| Reads Mapped Confidently to Intergenic Regions | 6.00% | 6.20% | 5.90% | 6.20% | 4.90% | 5.20% | 4.90% | 4.70% | 4.50% | 3.80% | 4.10% | 4.50% |
| Reads Mapped Confidently to Intronic Regions | 28.00% | 27.90% | 28.60% | 30.00% | 26.30% | 25.30% | 25.60% | 22.90% | 23.60% | 22.60% | 18.80% | 26.20% |
| Reads Mapped Confidently to Exonic Regions | 59.80% | 60.20% | 59.00% | 57.20% | 63.20% | 64.30% | 63.70% | 66.70% | 66.40% | 68.60% | 71.90% | 64.40% |
| Reads Mapped Confidently to Transcriptome | 56.20% | 56.50% | 55.20% | 53.40% | 59.70% | 60.60% | 59.80% | 62.90% | 62.80% | 65.60% | 68.60% | 61.30% |
| Reads Mapped Antisense to Gene | 1.50% | 1.60% | 1.80% | 1.70% | 1.50% | 1.50% | 1.80% | 1.60% | 1.50% | 0.90% | 1.10% | 1.10% |

| | | | | | | | | | | | | |
|---|---|---|---|---|---|---|---|---|---|---|---|---|
| **Fraction Reads in Cells** | 86.40% | 85.70% | 82.50% | 83.90% | 88.40% | 94.10% | 90.90% | 91.80% | 91.90% | 91.90% | 93.20% | 87.70% |
| **Total Genes Detected** | 25,924 | 25,469 | 25,746 | 26,366 | 25,710 | 26,445 | 26,434 | 26,274 | 26,389 | 24,967 | 24,826 | 24,950 |
| **Median UMI Counts per Cell** | 6,970 | 6,923 | 6,528 | 6,507 | 6,980 | 11,706 | 10,698 | 11,874 | 11,059 | 8,699 | 5,449 | 8,819 |

**Table S2. Deconvolution of shared single cell pools and doublet detection.**

| Pool | Pooled Individuals | Detected Individuals | Scrublet minimum gene variability percentage | Singlets | Doublets |
|---|---|---|---|---|---|
| 1 | 8 | 8 | 80 | 12,891 | 1,472 |
| 2 | 8 | 6 | 80 | 9,896 | 2,008 |
| 3 | 8 | 8 | 80 | 11,825 | 2,398 |
| 4 | 8 | 8 | 80 | 12,559 | 1,876 |
| 5 | 8 | 8 | 80 | 12,588 | 1,855 |
| 6 | 8 | 8 | 80 | 11,924 | 1,699 |
| 7 | 7 | 7 | 80 | 12,419 | 2,270 |
| 8 | 6 | 6 | 80 | 4,967 | 2,498 |
| 9 | 7 | 7 | 80 | 12,013 | 1,862 |
| 10 | 7 | 8 | 80 | 12,843 | 2,849 |
| 11 | 7 | 6 | 80 | 6,495 | 1,410 |
| 12 | 6 | 6 | 80 | 9,539 | 4,602 |
| | 88 | 86 | 90 | 129,959 | 26,799 |

**Table S3. Summary of cell type and donor assignments.**

| Reference | GA | Control | Total | Subpopulation |
|---|---|---|---|---|
| Common_4 | 32,254 | 24,027 | 56,281 | RPE1 |
| Common_6 | 14,620 | 16,780 | 31,400 | RPE2 |
| Common_3 | 14,501 | 5,851 | 20,352 | RPE3 |
| Common_1 | 4,329 | 2,113 | 6,442 | RPE4 |
| Control_3 | 2,977 | 3,159 | 6,136 | ProgenitorRPE |
| Common_2 | 2,269 | 1,396 | 3,665 | RPE5 |
| Control_0 | 2,211 | 1,172 | 3,383 | RPE6 |
| Control_4 | 186 | 98 | 284 | RPE7 |
| Control_2 | 221 | 41 | 262 | RPE8 |
| Aged_5 | 87 | 36 | 123 | RPE9 |
| Control_5 | 34 | 34 | 68 | RPE10 |
| Aged_1 | 33 | 21 | 54 | RPE11 |
| Control_1 | 8 | 12 | 20 | RPE12 |
| Aged_0 | 3 | 0 | 3 | RPE13 |
| Common_5 | 1 | 2 | 3 | RPE14 |
| Aged_2 | 1 | 0 | 1 | RPE15 |
| Aged_3 | 0 | 1 | 1 | RPE16 |
| | 73,735 | 54,743 | 128,478 | |

GA: geographic atrophy.

**Table S4. Post-hoc comparisons of cell subpopulation proportions between conditions**

| Subpopulation | Residuals (GA) | Adjusted P-values (GA) | Residuals (Control) | Adjusted P-values (Control) |
|---|---|---|---|---|
| ProgenitorRPE | -1.43E+01 | 0.00E+00 | 1.43E+01 | 0.00E+00 |
| RPE1 | -5.43E-03 | 1.00E+00 | 5.43E-03 | 1.00E+00 |
| RPE2 | -4.43E+01 | 0.00E+00 | 4.43E+01 | 0.00E+00 |
| RPE3 | 4.39E+01 | 0.00E+00 | -4.39E+01 | 0.00E+00 |
| RPE4 | 1.65E+01 | 0.00E+00 | -1.65E+01 | 0.00E+00 |
| RPE5 | 5.71E+00 | 1.60E-07 | -5.71E+00 | 1.60E-07 |
| RPE6 | 9.59E+00 | 0.00E+00 | -9.59E+00 | 0.00E+00 |

**Table S5. Statistical significance of differences between average cell subpopulation proportions**

| Subpopulation | Baseline Prop - Frequency | Prop Mean - GA | Prop Mean - Control | PropRatio | Tstatistic | P.Value | FDR |
|---|---|---|---|---|---|---|---|
| ProgenitorRPE | 0.048 | 0.036 | 0.053 | 0.681 | -1.510 | 0.135 | 0.207 |
| RPE1 | 0.441 | 0.452 | 0.472 | 0.957 | -0.481 | 0.632 | 0.632 |
| RPE2 | 0.246 | 0.176 | 0.284 | 0.619 | -3.095 | 0.003 | 0.019 |
| RPE3 | 0.159 | 0.215 | 0.112 | 1.928 | 2.427 | 0.017 | 0.061 |
| RPE4 | 0.050 | 0.057 | 0.037 | 1.536 | 2.024 | 0.046 | 0.108 |
| RPE5 | 0.029 | 0.035 | 0.025 | 1.430 | 1.462 | 0.148 | 0.207 |
| RPE6 | 0.027 | 0.029 | 0.018 | 1.589 | 1.234 | 0.221 | 0.258 |

**Table S6. Lead cis-eQTL and interactions with genotype and disease in RPE subpopulations**
See additional File

**Table S7. Common cis-eQTL in RPE1 and RPE2**
See additional File
